# The Genetic Basis of Bacterial Adaptation to Hosts

**DOI:** 10.1101/2025.07.19.665706

**Authors:** Ofir Segev, Alexander Martin Geller, Cristina Bez, Peihan Wang, Avital Akerman-Arad, Lusine Nazaretyan, Fei Chen, Vittorio Venturi, Asaf Levy

**Affiliations:** The Department of Plant Pathology and Microbiology, The Institute of Environmental Science, The Robert H. Smith Faculty of Agriculture, Food, and Environment, The Hebrew University of Jerusalem, Rehovot, Israel; The Bacteriology Group of the International Centre for Genetic Engineering and Biotechnology (ICGEB), Trieste, Italy; CAS Key Laboratory of Genome Sciences and Information, Beijing Institute of Genomics, Chinese Academy of Sciences and China National Center for Bioinformation, Beijing, China; Berlin Institute of Health at Charité-Universitätsmedizin Berlin, Berlin, Germany; African Genome Center, University Mohammed VI Polytechnic (UM6P), Ben Guerir, Morocco

## Abstract

Microbes colonize and interact with diverse multicellular hosts using specialized genes, many of which remain unidentified. Better understanding of host-associated gene functions is a key aspect of microbial ecology. We utilized a large-scale comparative genomics approach to identify and characterize host-associated functions by comparing 72,079 high-quality bacterial genomes from host and non-host environments using five enrichment tests for high accuracy. We uncovered over 3,000 protein domains, 16,000 AlphaFold protein clusters, and 1,500 operons enriched in host-associated bacteria. Additionally, we identified proteins and domains that are enriched in animal- or plant-associated bacteria. These include new functions such as mercury detoxification in hosts and animals in particular, and numerous proteins and domains of unknown function. We validated our results by genetically disrupting five poorly annotated host-associated genes in plant-associated bacteria, resulting in a substantial reduction in rice root colonization. One of the new colonization factors strongly affected bacterial motility and resistance of oxidative stress. Our findings, presented in a new database, GOTHAM DB, reveal the genetic basis of bacterial host association, including new functions underlying host-microbe interactions, and advance our understanding of microbial evolution.

## Main

Bacteria dwell in virtually every habitat on Earth through their diversity and adaptability. They engage in complex ecological interactions with their surrounding environment and other organisms, with host-microbe relationships often exerting significant positive or negative effects on the fitness of multicellular eukaryotes. For example, the microbiome plays a vital role in many aspects of the host’s life, from aiding in digestion and metabolism^1^, supporting and developing a proper immune system^2–5^, promoting host development^6^, actively protecting against pathogen invasion by secreting antimicrobials^7,8^, affecting host behavior^9^, or simply by occupying niches and competing for nutrients^10,11^. Additionally, beneficial microbes provide essential nutrients to their host, such as vitamin K to humans^12^, together with nitrogen, iron and phosphorus to plants^13,14^. On the other hand, pathogens have developed various strategies to colonize hosts, driving major public health and agricultural challenges^15,16^.

The molecular foundation of host-microbe interactions lies in bacterial genes that have co-evolved with their hosts, enabling microbial survival and proliferation in specialized niches^17^. Numerous studies have explored these actuators, but a comprehensive understanding has yet to be established. Traditional Tn-seq studies, while powerful for identifying fitness-essential microbial genes through mutant libraries^18–22^, are constrained to model organisms with established genetic tools and measurable phenotypes.

High-throughput sequencing has opened a broader avenue through comparative genomics, enabling researchers to identify host-associated genes by comparing bacterial genomes from eukaryotic hosts with their environmental counterparts. This approach has been used in PathFams for identifying Pfams enriched in pathogen genomes^23^ and in bacLIFE to detect lifestyle-associated genes^24^. While these approaches are valid, they often overlook the complexities of bacterial genome-wide association studies (GWAS), potentially leading to false positives. The clonal nature of bacterial growth results in many nearly identical genomes, violating the random sampling assumption of standard statistical enrichment methods. Population structure can bias results by correlating genetic variants with both the phenotype and the population structure itself. Using statistical tools that account for population structure helps to better clarify gene-trait associations^25–30^.

Our previous study comparing plant-associated and non-plant-associated bacterial genomes identified thousands of putative plant-associated protein orthologs through sequence-based clustering^31^. Recent clustering of 214 million AlphaFold-predicted proteins using FoldSeek^32^ formed 2.3 million structural clusters (AFCs), compared to 52.3 million sequence-based clusters^33^. This highlights the limitations of sequence-based clustering in enrichment analyses, though the use of protein 3D structures in comparative genomics is still in its infancy.

Here, we systematically identified host-associated bacterial genes via comprehensive analysis of 72,079 high quality bacterial genomes, classified as host-associated or environmental based on isolation sites. The comparative genomics analysis was performed at the genus and family levels using both protein domains and AlphaFold-based protein structural clusters (AFCs). Thousands of domains and proteins were independently enriched in multiple host-associated taxa. In addition, we identified predicted operons and pathways enriched in host-associated bacteria. By employing multiple phylogeny-aware statistical tests, we reduced potential biases in each method. Further comparison between animal- and plant-associated bacterial genomes revealed kingdom-specific host-association patterns. One surprising finding is that mercury resistance and certain transposons are enriched in hosts in general, and in animals in particular. Thousands of host-associated proteins have unidentified functions, which are over one-third of all enriched host-associated proteins. Importantly, we experimentally validated our approach by demonstrating that deletion of newly identified host-associated genes of unknown function significantly impaired rice root colonization of different bacterial strains. One of these genes is involved in motility and resistance to reactive oxygen species. Our findings are accessible through a new database, GOTHAM DB (https://ngdc.cncb.ac.cn/GeneAdaptionDB/), and provide a comprehensive genomic framework that advances our understanding of bacterial host adaptation. This computational approach reveals new proteins and domains involved in host adaptation, potentially improving our ability to engineer beneficial microbes, and identify new virulence functions. Illuminating the genetic determinants of host association provides an essential foundation for deciphering and manipulating bacterial-host interactions across diverse biological systems.

## Results

### Establishing a Database of Bacterial Isolates Labeled by Their Isolation Source

A total of 931,343 genomes of bacterial isolates were retrieved from various sources^34–43^. These genomes underwent stringent quality filtering, retaining only those with an N50 above 50,000 bp, completeness exceeding 95%, and contamination below 5% (n=744,298). Based on isolation source metadata, the filtered high quality genomes were labeled as host-associated or environmental. Due to the large dataset size, we manually labeled metadata for 70% of the genomes, which were then used to train a machine learning classifier for the remaining samples. This process was streamlined by the fact that many genomes shared identical metadata fields (e.g., host: “*Homo sapiens*”), allowing efficient annotation through bulk assignment rather than genome-by-genome labeling. Genomes lacking sufficient metadata or with ambiguous isolation sources (e.g., food or man-made facilities) were excluded from the database. Genomes with detailed metadata were further labeled with host type information, identifying them as either animal- or plant-associated. To minimize redundancy, the genomes were clustered based on sequence similarity^44^, and a representative genome from each cluster was randomly selected for the statistical analysis. This process was repeated independently three times to minimize bias from randomly selected genomes. Finally, to ensure consistency in taxonomy names across different sources, all genomes were assigned taxonomy classifications according to the GTDB^45^.

The final database contained 72,079 non-redundant, high quality genomes of bacterial isolates (Fig. 1a, Supplementary Table 1). This database exhibits remarkable diversity, encompassing genomes distributed across nine major bacterial phyla. Approximately 82% of the genomes were labeled as host-associated (Fig. 1b), a plausible proportion given the high prevalence of sequenced bacterial genomes isolated from humans^43^. Despite this, 114 genera and 66 families displayed a sufficient diversity of “host-associated” vs. “environmental” labels, facilitating meaningful enrichment analysis based on entropy calculation (Materials and Methods).

**Figure 1:**
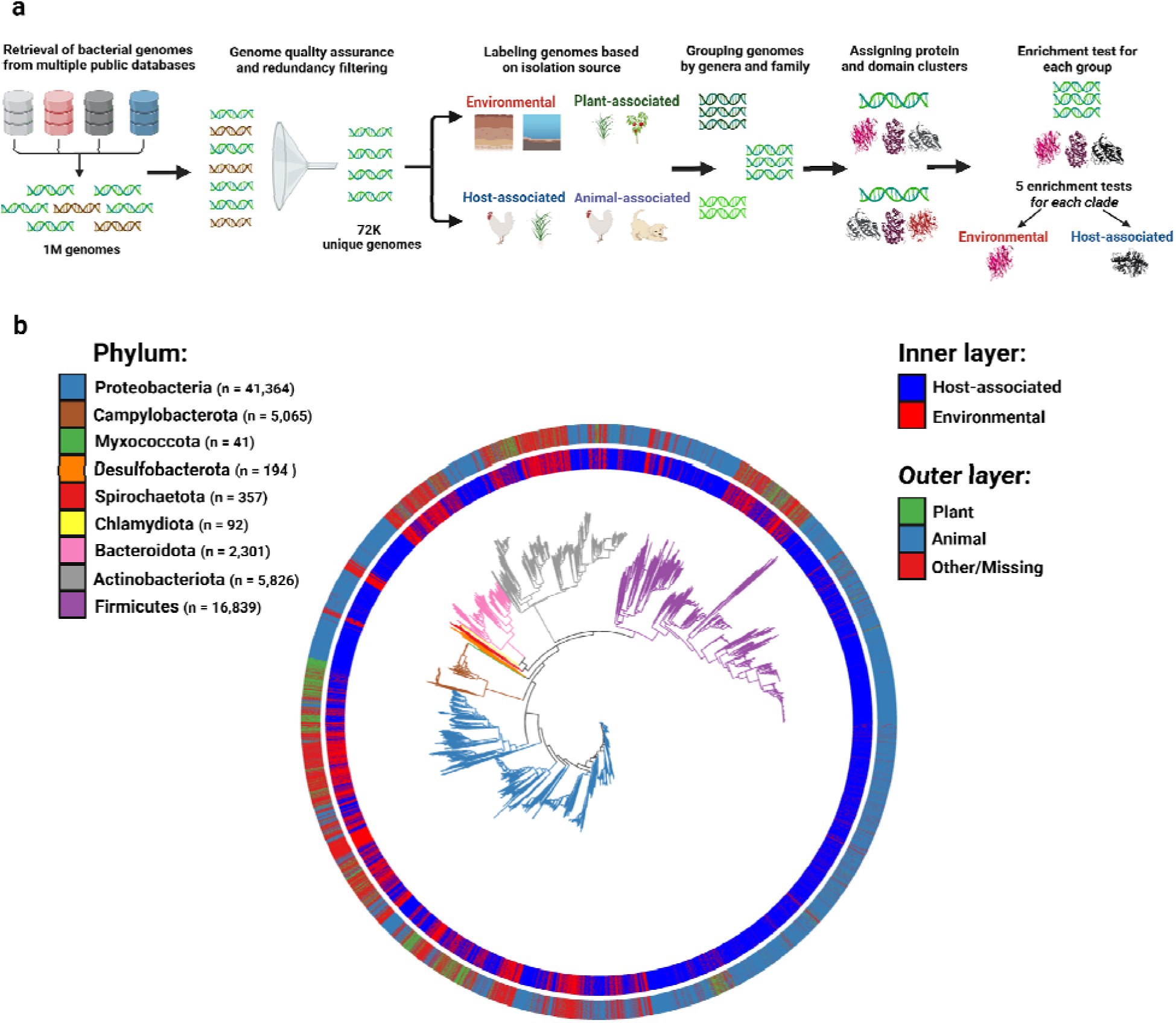
Comparative genomics method and resulting genome database. a. A Pipeline for Database Creation. Genome sequences (n=931,343) were retrieved from multiple repositories^34–43^. These sequences were filtered to ensure high quality and completeness. To eliminate redundancy, genomes were clustered based on nucleotide sequence similarity, with a representative genome selected from each cluster. The remaining high-quality, non-redundant genomes were labeled according to their isolation source which was based on the associated metadata. The genomes were further grouped by genus and family for a clade-specific comparison, and were further annotated with the protein clusters (AFCs) and protein domain clusters (Pfams) they encode. Finally, each AFC/Pfam and clade was subjected to five enrichment test to correlate it with a niche (host/environment and later also animal/plant). **b. A phylogenetic tree displaying microbial genomes meeting the established thresholds.** Colored branches denote distinct phyla. The tree is further annotated with layers indicating specific labels assigned to each genome. The inner layer denotes whether the genome is classified as host-associated, while the outer layer indicates the host identity, whether plant, animal, or neither.

For each genome in the database, all encoded genes were annotated for protein domains using the Pfam database^46^. To identify host-associated protein domains, genomes were grouped by genera and families. Each taxonomic clade was subjected to five distinct enrichment tests, comparing Pfam domain presence between host-associated and environmental genomes. Every test employs a unique methodology to assess Pfam enrichment, resulting in a major variation in their ability to accurately identify distinct enrichment patterns (Supplementary Fig. 1a). Scoary^25^, Pyseer^26^, Evolink^27^ (presence-absence version), and Fisher exact test assess the presence or absence of a protein domain, while Evolink^27^ (counts version) utilizes domain copy-number data. Furthermore, all tests, except for Fisher, incorporate the distribution of the Pfam domain within the phylogenetic tree as part of their calculation. As a result, the number of Pfam domains identified as enriched varies among the tests (Supplementary Fig. 1b). Given that lenient tests may yield more false positives while stringent tests might miss true positives, no single test can be considered superior in our view. The advantages and limitations of each method are discussed in the Materials and Methods section. To evaluate the significance of Pfam domain enrichment, we assigned a score ranging from 0 to 5 based on the number of tests that identified the Pfam domain as significantly host-associated. Pfam domains identified as enriched by multiple tests were considered to be more reliable in further analysis.

### Enrichment of Host-associated Pfam Domains Across Diverse Taxonomic Groups

Each genus was analyzed separately, and the results were then combined, revealing a total of 3,280 distinct Pfam domains enriched in host-associated bacteria across 114 genera, with 1,162 domains supported by two or more enrichment tests (Supplementary Table 2). Pfam domains enriched by at least one test across multiple phylogenetically distinct clades are of particular interest, as they are more likely to play an adaptive role in the host niche. In our analysis, 16% of the enriched host-associated Pfam domains (n=526) were identified in at least three genera. A representative subset of 30 highly enriched Pfam domains across multiple genera is shown in Fig. 2a.

**Figure 2:**
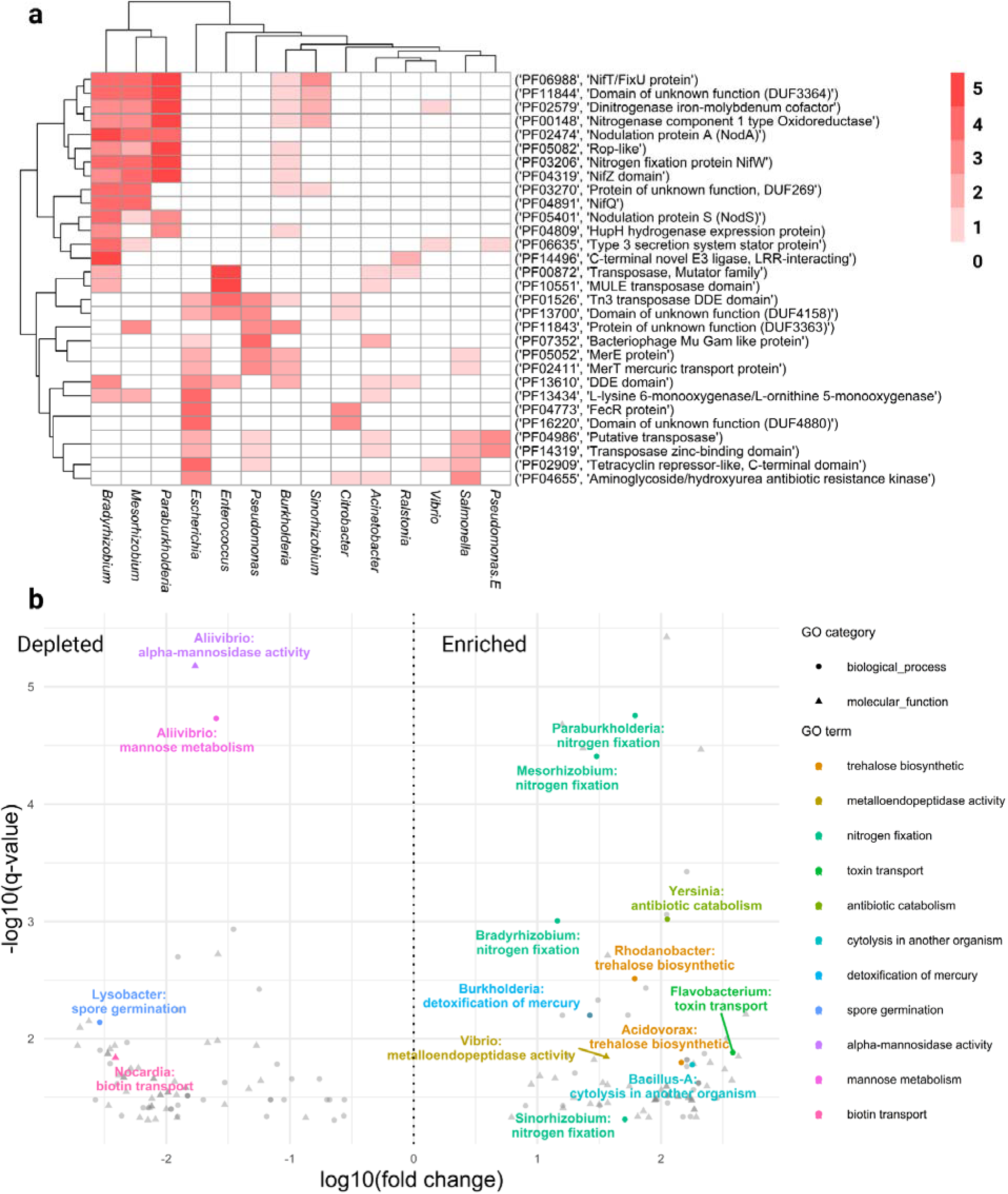
Enrichment of Host-associated Pfam Domains at the Genus level. a. Heatmap showing the enrichment of host-associated Pfam domains across 14 bacterial genera. Each row represents a Pfam domain (Pfam ID and description), and columns correspond to genera. Red shading indicates the number of tests identifying these domains as significantly enriched in host-associated bacteria within a clade, with white indicating no enrichment.**b. Volcano plot highlighting highly enriched host-associated functions based on Pfam domains.** The y-axis represents the significance of enrichment (-log10 FDR-corrected q-value), while the x-axis shows the fold change (log10 fold change) for each mapped GO term. Colors highlight specific key functions. Functions located to the right of the dotted line are enriched in host-associated bacteria, while those on the left are depleted. To improve readability, “trehalose biosynthetic process” was abbreviated to “trehalose biosynthesis”, “antibiotic catabolic process” to “antibiotic catabolism”, “detoxification of mercury ion” to “detoxification of mercury”, “biotin transmembrane transporter activity” to “biotin transport” and “mannose metabolic process” to “mannose metabolism”.

The most prominently enriched domains are linked to classic rhizobial functions: root nodulation and nitrogen fixation. These domains, including nifT, nifW, nifZ, nifQ, DUF3364, NodA, and NodS, were consistently significant in *Sinorhizobium*, *Mesorhizobium*, *Bradyrhizobium*, *Burkholderia*, and *Paraburkholderia*, playing key roles in rhizobia-legume interactions^47–49^. Additional host-associated domains include the “C-terminal novel E3 ligase, LRR-interacting” (PF14496), that is linked to bacterial virulence^50^ and is enriched in *Ralstonia* and *Bradyrhizobium*. We also identified the “Type 3 secretion system stator protein” domain (PF06635), which regulates ATPase of T3SSs that inject bacterial effectors into eukaryotic cells^51^, as enriched in *Vibrio*, *Pseudomonas-E*, *Bradyrhizobium*, and *Mesorhizobium*. Notably, additional virulence factors, such as Yersinia/Haemophilus virulence surface antigen (PF03543) and IucA/IucC family domain (PF04183), and T3SS components, such as the periplasmic domain of Yop proteins (PF16693) and a cytoplasmic component of the T3SS needle (PF08988), were identified (Supplementary Table 2).

Various mobile elements, such as the “MULE transposase domain” and “Tn3 transposase DDE domain” were enriched in host-associated bacteria, suggesting their potential role in transferring host-related functions. Interestingly, Pfams related to antibiotic and mercury resistance, such as “aminoglycoside/hydroxyurea antibiotic resistance kinase” (PF04655), merE, and merT^52^, were independently enriched across host-associated *Escherichia*, *Burkholderia*, *Pseudomonas*, *Salmonella*, and others. Despite the unexpected enrichment, mercury detoxification has also been observed in bacteria isolated from primate feces^53^. We propose that its prevalence in predominantly human-associated genera may reflect adaptation to elevated mercury levels in humans, potentially originating from dental fillings^54^, contaminated fish^55^, or other pollutants. Notably, multiple domains of unknown function, such as DUF3363 and DUF4880, were highly enriched, suggesting novel potential roles in host association. Conversely, the shared domains enriched in environmental bacteria displayed no clear pattern of widespread enrichment. (Supplementary Fig. 3).

Given the vast number of enriched host-associated Pfam domains, we aimed to characterize broader patterns of host-association by correlating these enriched domains with Gene Ontology (GO) functional terms. To achieve this, we focused on Pfam domains that were enriched in more than one test within specific genera. For each genus, we calculated the GO term enrichment by counting the occurrences of GO terms associated with enriched Pfam domains and normalizing by their overall distribution across all Pfam domains encoded by the genus (Fig. 2b, Supplementary Table 3). Our analysis revealed a significant representation of processes related to host adaptation, such as nitrogen fixation by bacteria in legume root nodules, and “cytolysis in another organism” in *Bacillus-A*. Moreover, trehalose biosynthesis, enriched in *Rhodanobacter* and *Acidovorax*, is proposed to aid host colonization by enhancing stress tolerance, modulating immune responses, and promoting pathogenicity^56^. Additionally, toxic substances degradation such as mercury detoxification and antibiotics catabolism were prominent in *Burkholderia* and *Yersinia*, respectively. In contrast, functions like spore germination were underrepresented in host-associated bacteria, suggesting a shift away from adaptation to extreme environments.

We applied the same statistical methodology to genomes grouped by family. We identified 4,357 Pfam domains enriched across 66 families, of which 1,045 were supported by two or more tests (Supplementary Fig. 4-5, Supplementary Table 2). Cellulase (PF00150), for example, was found as host-associated in six families and as an environmental domain in three other families. GO analysis of these enriched Pfams revealed that carbohydrate metabolism and import, particularly xylan catabolism—a key component of plant cell walls^57^, are linked to host association in certain families. Nitrogen fixation also remained a prominent feature of host-associated bacteria (Supplementary Fig. 6, Supplementary Table 3).

### Enrichment of Host-associated AlphaFold Clusters

To complement the host-associated Pfam domain enrichment analysis, we investigated enrichment at the full-protein level. This additional dimension could not only strengthen the validity of already enriched protein domains but also reveal the enrichment of multidomain proteins or proteins that could not be mapped to any Pfam domains.

To achieve this, we first clustered all proteins encoded by the bacterial genomes in our database using a structure-based approach. Leveraging the existing AlphaFold clusters (AFC) database^33^, we matched each protein sequence to a closely related UniProt protein sequence, thereby linking it to its corresponding AlphaFold cluster (Supplementary Table 4). Using structure-based instead of sequence-based clusters maximizes clustering completeness as proteins with different sequences but similar structures, which often indicate shared functions, are grouped into the same cluster.

Using the AFCs mapping of the genes in our database, host-associated proteins were identified by applying the five previously mentioned enrichment tests. This resulted in the enrichment of 16,446 distinct AFCs in host-associated bacteria across 114 genera, with 3,469 passing two or more tests (Supplementary Table 5). The greater number of enriched AFCs compared to Pfams reflects the richer functional information from AFC clusters (n=800,000), resulting in a dataset 40 times larger than Pfam.

A representative subset of enriched and depleted host-associated AFCs across multiple genera is presented (Fig. 3a, Supplementary Fig. 7). Notably, a substantial proportion of the most widespread host-associated AFCs are uncharacterized proteins (Fig. 3a, Supplementary Table 5). In fact, over one-third of all enriched host-associated AFCs are uncharacterized proteins, highlighting the many host-associated genes that remain unexplored. However, some enriched AFCs are linked to known host interactions, consistent with Pfam-based results, including nitrogenase activity essential for nitrogen fixation, T3SS proteins, and TonB-dependent receptors involved in the uptake of siderophores, vitamin B12, and carbohydrates^58^, all of which are present in the host environment. Additionally, conjugal transfer proteins were enriched in *Pseudomonas* and *Burkholderia*, suggesting their role in transferring genetic elements that may facilitate host adaptation in neighboring bacteria.

**Figure 3:**
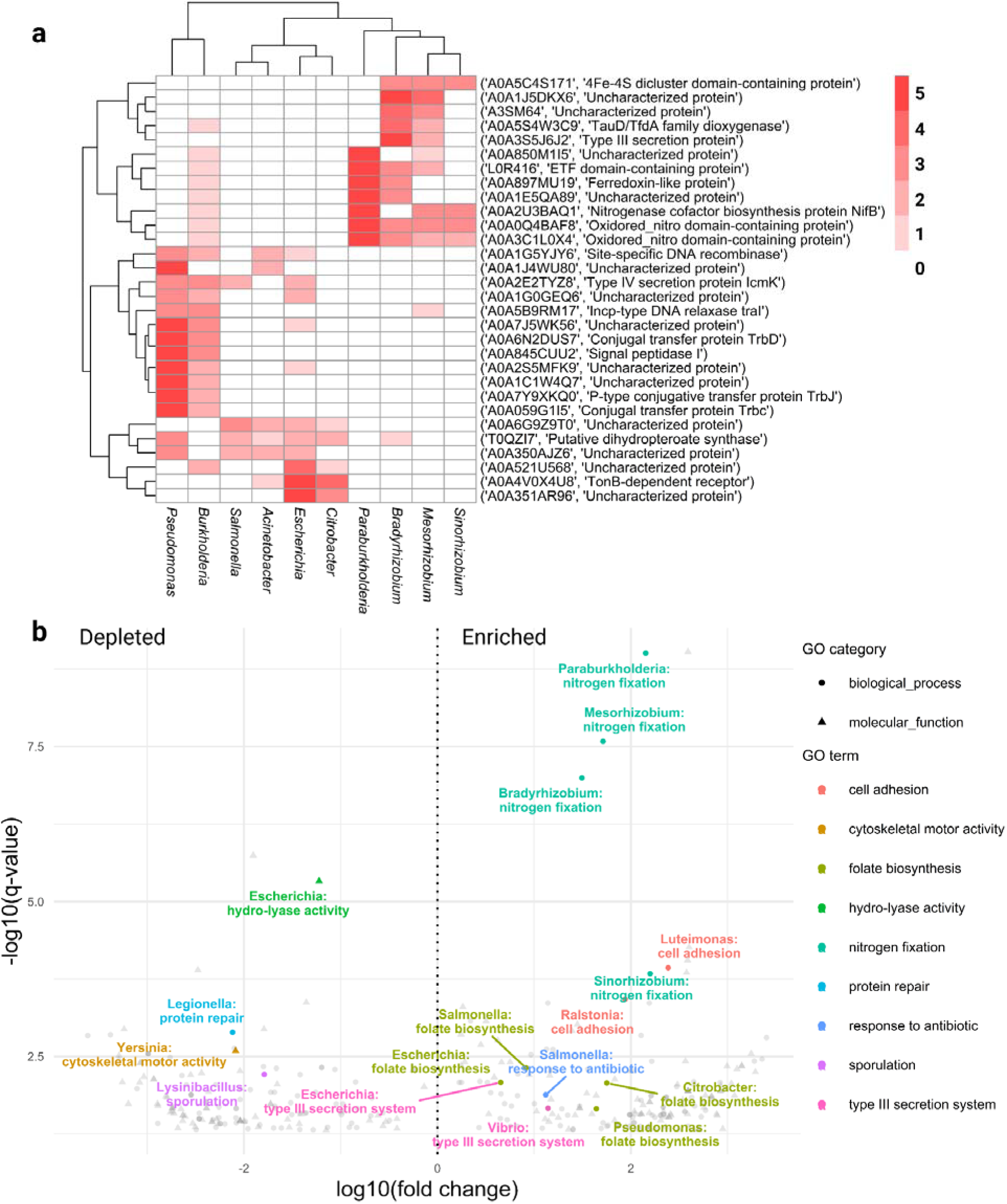
Enrichment of Host-associated AlphaFold clusters at the Genus level. a. Heatmap showing the enrichment of host-associated AlphaFold clusters across 10 bacterial genera. Each row represents an AlphaFold cluster (Cluster ID and description), and columns correspond to genera. Red shading indicates the number of tests identifying these domains as significantly enriched in host-associated bacteria within a clade, with white indicating no enrichment. **b. Volcano plot highlighting highly enriched host-associated functions based on AlphaFold clusters.** The y-axis represents the significance of enrichment (-log10 FDR-corrected q-value), while the x-axis shows the fold change (log10 fold change) for each mapped GO term. Colors highlight specific key functions. Functions located to the right of the dotted line are enriched in host-associated bacteria, while those on the left are depleted. To improve readability, “folic acid-containing compound biosynthetic process” was abbreviated to “folate biosynthesis” and “protein secretion by the type III secretion system” to “type III secretion system”.

To gain insights into the functions of host-associated AFCs, we mapped all AFCs to GO terms using information from the UniProt database^59^. We then applied a similar approach as used for Pfam domains to identify enriched GO terms (Fig. 3b, Supplementary Table 6).

The mapped results mirrored patterns observed with Pfam domains, highlighting nitrogen fixation in host-associated bacteria across four rhizobial genera. T3SS was enriched in *Vibrio* and *Escherichia*, while functions that may aid in host adaptation like cell adhesion (*Luteimonas, Ralstonia*) and antibiotic response (*Salmonella*) were also prominent. Interestingly, the biosynthesis of folic acid-containing compounds was enriched in four distinct genera, suggesting a potential role in host adaptation. Folates play a crucial role in host health, as they are essential for DNA synthesis and epigenetic regulation^60^. Since mammalian cells cannot produce folates, they rely on dietary sources and intestinal microbiota for their supply^61^. While some gastrointestinal strains can synthesize folates de novo, many rely on p-aminobenzoic acid (pABA), a precursor obtained either from the host’s diet or other gut microbes^60^. This abundance of pABA in the gut likely drives host-associated bacteria to adapt by producing more folate compounds, benefiting both the bacteria and the host. On the other hand, pathogens also benefit from folate acid metabolism which have been demonstrated to play a role in virulence^62^. Moreover, antifolate antibiotics are commonly used to combat bacterial infections^63^.

Enriched host-associated AFC results at the family level include 30,985 AFCs across 66 families, of which 4,528 passed two or more tests (Supplementary Fig. 8-9, Supplementary Table 5). When mapped to GO terms, those enriched AFCs revealed additional host-associated traits, including host interaction, symbiont entry, and xylan catabolism (Supplementary Fig. 10, Supplementary Table 6), highlighting clear adaptation to the host niche. Depleted AFCs included arsenic resistance in Enterobacteriaceae (Supplementary Fig. 9) and oxidative stress response in three families (Supplementary Fig. 10). Notably, nearly 70% of genus-level enriched AFCs were also enriched at the family level, indicating consistency between the two analyses.

### Genomic Organization of Enriched Host-associated Functions

Some host-associated functions are complex and encoded by operons composed of enriched AFCs and Pfam domains. Prokaryotic operon structure allows bacteria to co-regulate expression of functionally related genes in a simple manner, and also facilitates the horizontal gene transfer of full functions/pathways^64^. To investigate whether the enriched results correspond to predicted functional operons, we scanned each genus for Pfams that repeatedly co-occur in close proximity across multiple host-associated genomes (Materials and Methods). This analysis identified 2,287 enriched Pfam domains organized into 1,491 gene clusters across 94 genera (Supplementary Table 7). We provide five examples of genomic contexts where enriched Pfam domains and their corresponding AFCs are likely co-expressed and function as operons (Fig. 4). In some of the cases the host-associated enrichment is within another genus.

**Figure 4:**
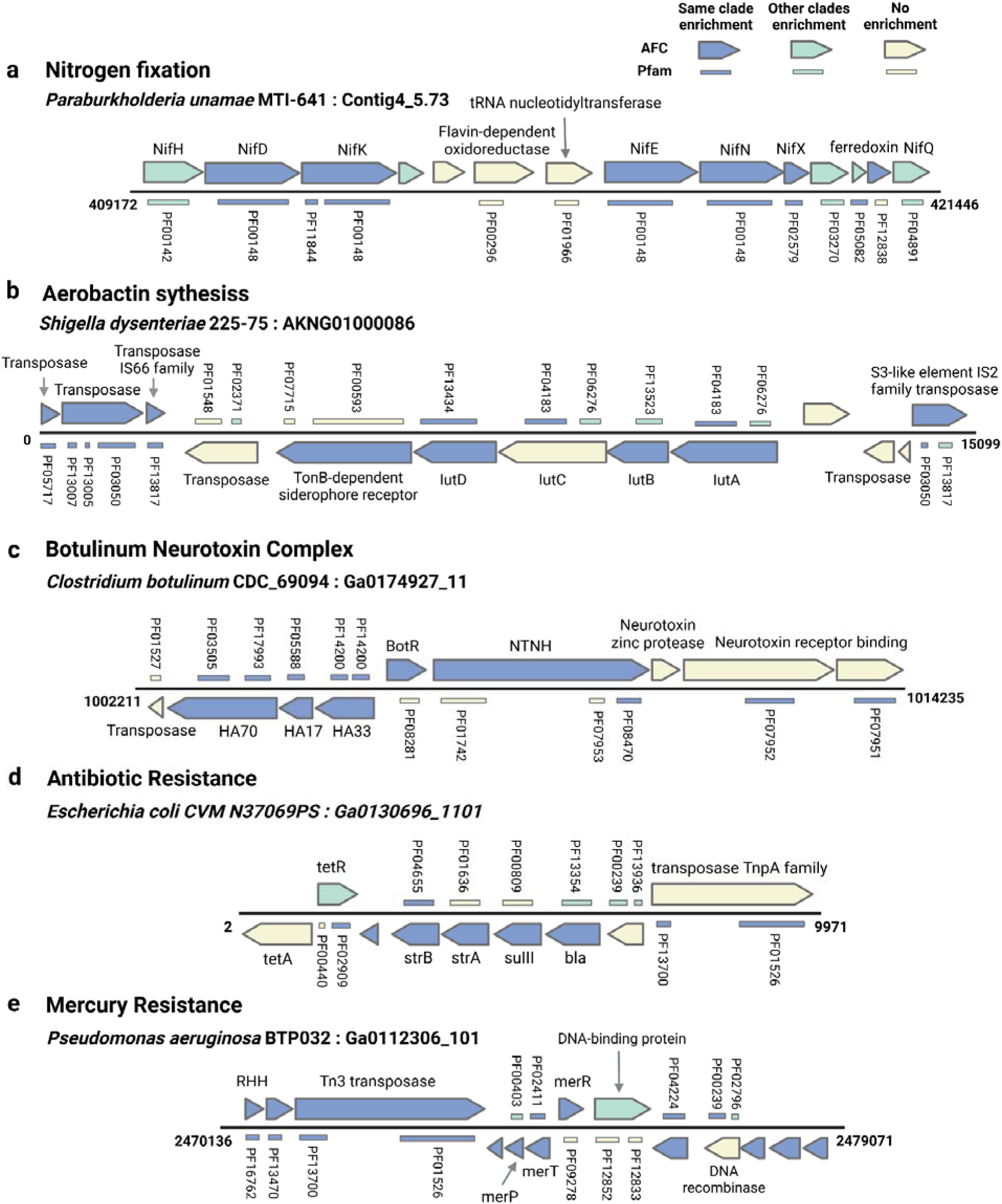
Functional operons and gene cassettes with enriched host-associated AFCs and Pfams. Each panel represents a functional operon found in the genome of a host-associated bacterium from our database. Genes highlighted in blue indicate enrichment within the same genus as the depicted strain, while teal indicates enrichment in other genera only. AFC enrichment is shown by the colored genes, and Pfam enrichment is indicated by the lines adjacent to those genes. **a.** Nitrogen fixation operon enriched in host-associated *Paraburkholderia*. b. Aerobactin biosynthesis operon enriched in host-associated *Shigella*. c. Botulinum neurotoxin complex enriched in host-associated *Clostridium*. d. Antibiotic resistance operon enriched in host-associated *Escherichia*. e. Mercury resistance operon enriched in host-associated *Pseudomonas aeruginosa*.

Operons encoding symbiosis functions, such as nitrogen fixation and nodulation, were identified, with most genes and domains enriched in host-associated bacteria (Fig. 4a, Supplementary Fig. 11), serving as a positive control for our approach. Additionally, operons for virulence factors were found, including the aerobactin iron transport and synthesis genes that are mapped within a pathogenicity island of *Shigella dysenteriae* (Fig. 4b). Aerobactin is a high-affinity iron chelator linked to increased virulence in enteric bacteria^65^. Another example is host-associated *Clostridium* (*Clostridium-F* by GTDB^45^), which showed significant enrichment in genes and domains related to the botulinum neurotoxin complex (Fig. 4c, Supplementary Fig. 12), responsible for both the expression of the botulinum toxin and its penetration into host cells^66^.

Operons conferring various resistances were also identified using this approach. A notable example is a functional operon in an *E. coli* strain that provides multiple antibiotic resistances, including *strA* and *strB*, encoding streptomycin-inactivating enzymes, *sulII*, conferring sulfonamide resistance, and *bla*, conferring beta-lactam resistance, all highly enriched across multiple genera (Fig. 4d). Similarly, the widespread enrichment of mercury resistance, initially surprising in the context of individual Pfam domains (Fig. 2a), was also identified as an enriched functional operon in the pathogen *Pseudomonas aeruginosa* (Fig. 4e) and other taxa, further reinforcing the evidence of mercury resistance adaptation among host-associated microbes.

Notably, mobile genetic elements adjacent to host-related functional genes were enriched alongside these genes (Fig. 4b,d-e). For example, PF13700 and PF01526, consistently enriched in multiple host-associated bacteria (Fig. 2b), are found in Tn transposases within antibiotic and mercury resistance operons (Fig. 4d-e). Furthermore, these functions may have been co-selected, aligning with reports that mercury resistance genes (*mer*) often travel with antibiotic resistance genes on mobile elements^67^.

### Computational Validation

To systematically validate our findings, we hypothesized that enriched Pfams and AFCs could effectively separate genomes within the same clade based on their isolation niches. We compared the ability to distinguish host-associated from environmental and plant versus animal bacteria using either all encoded Pfams/AFCs versus only those identified as enriched. Logistic regression modeling assessed separation improvement (Supplementary Figures 13–16, Supplementary Table 8). Across all combinations of taxonomic levels (genus and family), niche comparisons (host vs. environmental and plant vs. animal as explained below), and protein classification systems (AFCs and Pfams), 94% (456 out of 486) of tested groups showed improved Akaike information criterion (AIC) separation while maintaining F1 score performance when using only enriched results. F1 score is the harmonic mean of precision and recall. For example, in Nocardioidaceae family (Actinomycetota phylum) we observed a clear genome separation and reduction in AIC between host-associated and environmental bacteria only when we use enriched Pfams (Figure 5a) or enriched AFCs (Figure 5b) and not all possible Pfams and AFCs. This computationally confirms that these specific protein families are genuinely associated with niche-specific adaptation and provides a framework for prioritizing taxa where enriched predictions are most informative for ecological classification (Supplementary Table 8).

**Figure 5.**
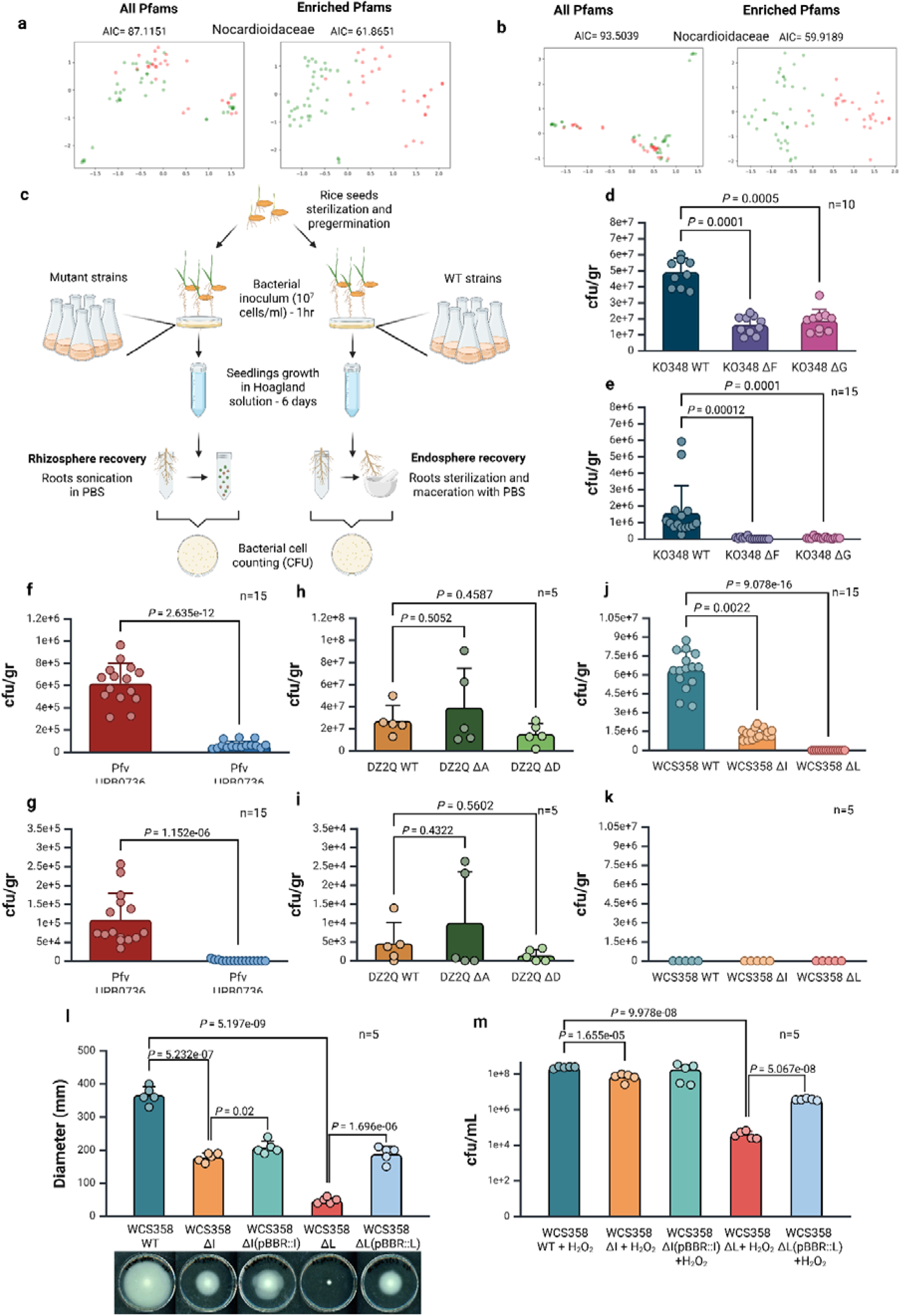
Computational and experimental validation of predicted host-associated genes. a-b. PCoA results of Nocardioidaceae labeled as animal-associated in red and plant-associated in green. On the left side are the genomes clustered by all genes mapped to AFC/Pfams, on the right side are the genomes clustered solely by enriched AFC/Pfams results. **a** PCoA results that show an improvement using the enriched Pfams results. **b** PCoA results that show an improvement using the enriched AFC results. **c, Experimental workflow showing the root colonization experiment.** Starting from the single inoculation of each mutant and the respective wild-type bacterial strain (107 cells/ml) onto pre-germinated rice seedlings. Rhizosphere recovery involved root sonication in PBS, while endosphere recovery required sterilization with ethanol (75% v/v) and hypochlorite (50% v/V) and maceration of roots in PBS. Serial dilutions of recovered cells were plated, and colony-forming units (CFUs) were counted after 24 hours. **d-e, Root colonization experiment of *Kosakonia oryzae* KO348**. Quantification of CFUs/gr of root in the rhizosphere (d) and endosphere (e) of seedlings inoculated with *Kosakonia oryzae* KO348 WT, KO348 ΔF, or KO348 ΔG. **f-g, Root colonization experiment of *Pseudomonas fuscovaginae* UPB0736**. Rhizosphere (f) and endosphere (g) colonization by *Pseudomonas fuscovaginae* Pfv UPB0736 WT and the mutant Pfv UPB0736 ΔB. **h-i, Root colonization experiment of *Dickeya zeae* D2Q2**. Rhizosphere (h) and endosphere (i) colonization by *Dickeya zeae* D2Q2 WT, D2Q2 ΔA, and D2Q2 ΔD. **j-k, Root colonization experiment of *Pseudomonas capeferrum* WCS358**. Rhizosphere (j) and endosphere (k) colonization by *Pseudomonas capeferrum* WCS358 WT, WCS358 ΔI, and WCS358 ΔL. **l, Swimming motility test of *Pseudomonas capeferrum* WCS358.** WT vs mutants WCS358 ΔI, WCS358 ΔL, complemented WCS358 ΔI(pBBR::I) and WCS358 ΔL(pBBR::L) on M9 solid medium (3% agar). **m, H2O2-kill assay of *Pseudomonas capeferrum* WCS358.** WT vs mutants WCS358 ΔI, WCS358 ΔL, complemented WCS358 ΔI(pBBR::I) and WCS358 ΔL(pBBR::L) under 2mM H2O2. Overnight cultures in Tryptic Soy broth (TSB) were inoculated into ME medium [20 g/L glucose, 12 g/L sodium citrate, 7 g/L (NH4)2SO4, 0.50 g/L K2HPO4·3H2O, 0.50 g/L MgSO4·7H2O, 0.04 g/L FeCl3·6H2O (0.15 mM Fe3+), 0.01 g/L MnSO4·H2O, and 0.15 g/L CaCl2, pH 7.5] and grown at 30°C for 12 h. Cells were then treated with 2 mM H2O2, and after 15 min of incubation, samples were diluted and plated on Tryptic soy broth agar (TSA), and incubated at 30°C for 16-18 h. Serial dilutions of recovered cells were plated, and cell survival was determined by counting CFU and is shown as the mean value ± standard error from five independent experiments. To assess statistical significance in **c-l**, Kruskal-Wallis and Dunn’s multiple comparison test was used. To assess statistical significance in **l-m**, One-way ANOVA and Dunnett multiple comparison test was used. All data are presented as means ± standard error (SD). The error bars indicate standard deviations.

### Genomic Knockout Mutants of Host-associated Genes Lead to Plant Colonization Impairment

To further validate our findings, we selected six Pfam domains that correspond to six AFCs, identified as enriched in host-associated bacteria and with no clear annotation for experimental validation. DUF3364 is an exception, as it was highly enriched but located within the *NifK* gene. Genome knockout mutations targeting seven genes in four strains (A,B,D,F,G,H,I), corresponding to the selected candidates, were constructed to assess their roles in host adaptation (Supplementary Table 9, Fig. 5c). The knockout mutant significantly impaired rice root colonization for 5 out of 7 mutants (Fig. 5d-k), while all strains displayed growth *in vitro* comparable to the wild-type strain (Supplementary Fig. 17,18). Therefore, we conclude that the deleted host-associated genes play a role *in planta*, in line with the notion that they are host-associated.

In *Kosakonia oryzae* KO348, ΔF and ΔG mutants exhibited a significant reduction in bacterial colonization of both the rhizosphere and endosphere compared to the WT (Fig. 5d-e). Similarly, *Pseudomonas fuscovaginae* UPB0736 ΔB mutants showed a substantial decrease in rhizosphere and endosphere colonization (Fig. 5f-g). In contrast, *Dickeya zeae* D2Q2 (a rice pathogen) mutants ΔA and ΔD, retained colonization ability (Fig. 5h-i). These observations suggest that the targeted genes in *Dickeya* may have strain-specific host functions or roles unrelated to root colonization. *Pseudomonas capeferrum* WCS358 is a well-studied rhizospheric strain^68^ that cannot colonize the endosphere (Fig. 5j-k). Notably, the deletion of genes *I* and *L* significantly reduced rhizosphere colonization, with the ΔL mutation eliminating it entirely (Fig. 5j). Overall, 5 of the 7 mutants showed reduced root colonization, supporting the hypothesis that the enriched host-associated proteins and domains contribute to bacterial adaptation to the host.

We further investigated whether the predicted host-associated candidate genes may play a role in bacterial movement towards and within the host. Motility driven by chemotaxis, facilitates migration to the rhizosphere, enabling attachment to plant roots, movement within roots, and the establishment of beneficial or pathogenic interactions^69,70^. Swimming assays revealed reduced motility in mutants *I* and *L* compared to the highly motile WT strain (Fig. 5l). Complementing the mutants in trans restored motility, confirming their involvement in bacterial motility (Fig. 5l). Notably, gene L appears to be essential for motility, as its deletion completely abolishes motility. Motility was shown before as a critical phenotype for root colonization^69–73^

To gain a better understanding of the functions of these genes, we examined their genetic context. In *P. capeferrum* WCS358, genes *I* and *L* were found adjacent to a gene encoding an enzyme with glutathionylspermidine synthase activity, containing PF03738, which was also enriched in host-associated *Pseudomonas-E (*including *P. capeferrum)* and other genera (Supplementary Fig. 19a). This reaction is crucial in pathogenic trypanosomatids for synthesizing the antioxidant trypanothione, which defends against reactive oxygen species (ROS)^74^ with homologs of this enzyme also present in *E. coli*^74^. Thus, we speculated that this gene cluster may function as a defense mechanism against ROS during bacterial invasion, as ROS are known to act as antimicrobial agents in the plant defense response^75,76^.

Interestingly, in *K. oryzae* KO348, gene F was located upstream within an operon that resembled the operon in *P. capeferrum* (Supplementary Fig. 19b). This operon contained the previously mentioned genes as well as *pspA (*A0A1E4K5N4, PF04012) and gene *F*, both of which were enriched in host-associated bacteria. *PspA* is a phage shock protein that stabilizes the bacterial cell membrane and protects against envelope stress^77^, further supporting its role in ROS defense.

To validate our hypothesis, we conducted an oxidative stress experiment, exposing ΔI and ΔL mutant strains of *P. capeferrum* WCS358 and the WT to HLOL (Fig. 5m). Remarkably, both knockout strains exhibited a significant reduction in bacterial survival compared to the WT, with ΔL presenting 10^4^ growth reduction and ΔI demonstrating a x3 growth reduction. The mutant strains harboring the wild type genes on a plasmid complemented the growth, underscoring the role of these genes in oxidative stress resistance. We conclude that gene I and L are new genes that contribute to existing bacterial mechanisms that combat oxidative stress to enable rice endophytic colonization^78^.

### Comparison Between Animal-Associated and Plant-Associated Bacteria

Animals and plants are the two major bacterial host kingdoms. Within the host-associated bacteria we could identify and label 53K and 5.5K animal-associated and plant-associated bacteria respectively (Fig. 1b), based on their isolation sites. We identified Pfam domains and AFCs enriched in animal-associated bacteria compared to plant-associated bacteria by applying all five enrichment tests on the genomic data with these additional labels. Dividing genomes at the genus level led to smaller clades and fewer significant findings, affecting the enrichment analysis. However, analyzing data at the family level provided more informative results across 35 families (Supplementary Table 10). These results include several widespread key functions, indicating adaptations specialized to specific host kingdoms (Supplementary Table 11). Figure 6 presents the findings based on enriched Pfam domains at the family level, with additional comparisons incorporating enriched AFCs, provided in Supplementary Figures 20 and 21.

**Figure 6:**
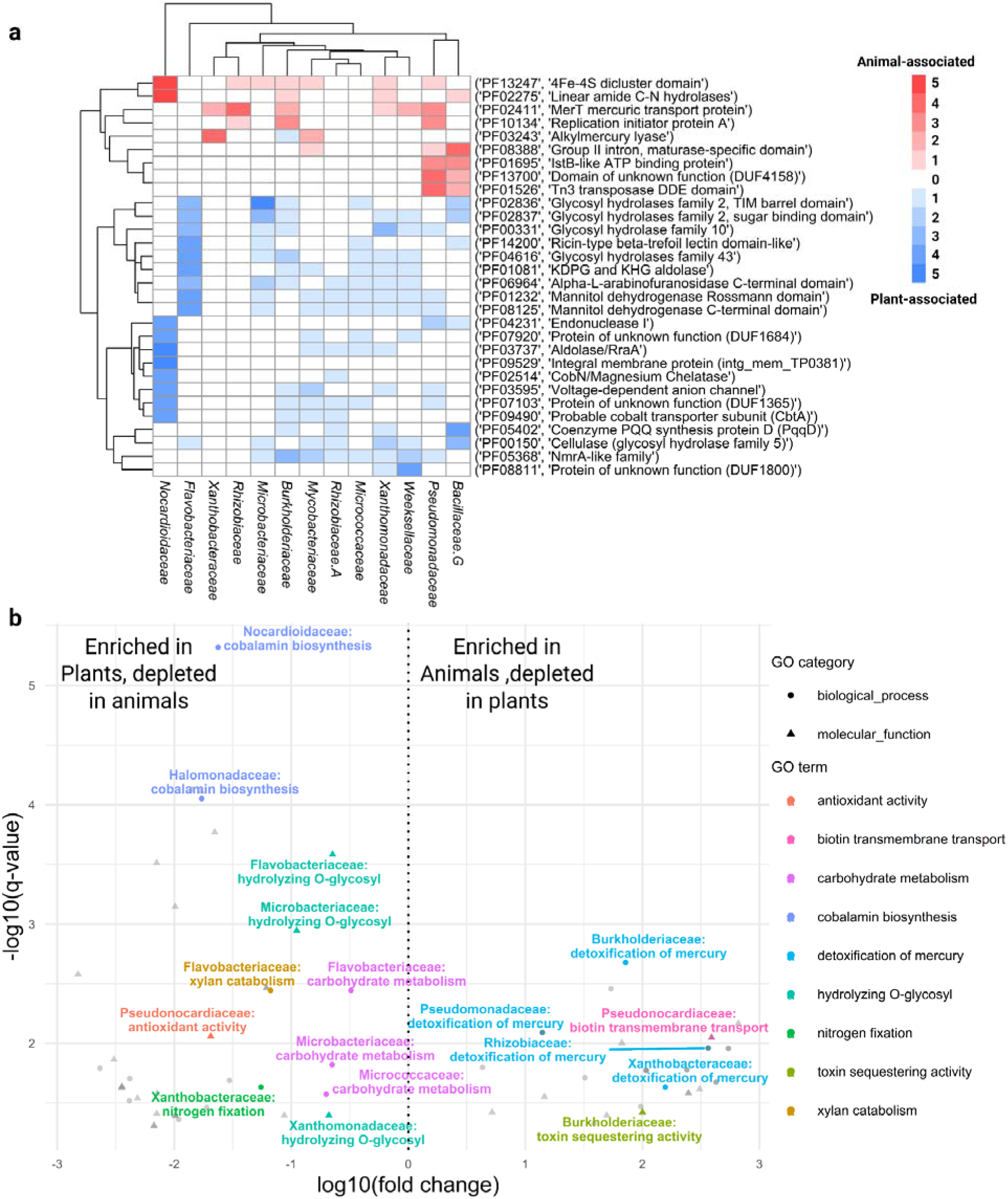
Enrichment of Animal-associated and Plant-associated Pfam Domains. a, Heatmap showing the enrichment of Animal-associated and Plant-associated Pfam domains across 13 bacterial families. Each row represents a Pfam domain, identified by its Pfam ID and description, while the columns correspond to different families. Red shading indicates the number of test that identified these domains as significantly enriched in animal-associated bacteria within a clade, and blue shading indicates enrichment in plant-associated bacteria. White areas denote no significant enrichment. **b, Volcano plot highlighting highly enriched animal-associated and plant-associated functions based on Pfam domains at the family level.** The y-axis represents the significance of enrichment (-log10 FDR-corrected q-value), while the x-axis shows the fold change (log10 fold change) for each mapped GO term. Colors highlight specific key functions. Functions located to the right of the dotted line are enriched in animal-associated bacteria, while those on the left enriched in plant-associated bacteria. To improve readability, “biotin transmembrane transporter activity” was abbreviated to “biotin transmembrane transport”, “hydrolase activity, hydrolyzing O-glycosyl compounds” to “hydrolyzing O-glycosyl”, “detoxification of mercury ion” to “detoxification of mercury”, “carbohydrate metabolic process” to “carbohydrate metabolism”, “cobalamin biosynthetic process” to “cobalamin biosynthesis” and “xylan catabolic process” to “xylan catabolism”.

Among the widespread animal-associated Pfams, merT, a domain found in the mercury resistance operon (Fig. 4e), was enriched across six families from Proteobacteria and Bacteroidetes phyla (Fig. 6a), alongside the functions “detoxification of mercury” prominent in four families (Fig. 6b) and “mercury ion transport” in Pseudomonadaceae (Supplementary Figure 21). The additional high enrichment of this function in general host-associated bacteria compared to environmental ones (Fig. 2, 4e), suggests that bacterial mercury resistance plays a crucial role in adapting specifically to animal hosts. Additionally, Tn3 transposase (PF01526) and DUF4158 (PF13700), which were observed within mobile elements in two of the previously discussed operons (Fig. 4d-e) and are likely involved in facilitating antibiotic and heavy metal resistance^79^, were also enriched in animal-associated bacteria (Fig. 6a), hinting at a specialization to animal hosts. The hemerythrin HHE cation binding domain (PF01814) responsible for oxygen transport in marine invertebrates^80^ was enriched in five animal-associated families from three phyla (Supplementary Table 10).

In plant-associated bacteria, nitrogen fixation and xylan catabolism are notably enriched (Fig. 6b), providing strong validation for identifying host-adaptive functions specialized to the plant kingdom. Additionally, many enriched widespread Pfam domains in plant-associated bacteria are involved in the hydrolysis of various sugars, including those abundant in plants such as cellulose (Fig. 6a), aligning with carbohydrate metabolism and the hydrolysis of O-glycosyl compounds, which are highly enriched across three families (Fig. 6b). This suggests that the utilization of plant carbohydrates is a key feature of bacterial adaptation to plant hosts, an observation that has been previously reported^41^. Notably, cobalamin (vitamin B12) biosynthesis shows the highest enrichment in plant-associated bacteria across Nocardioidaceae and Halomonadaceae (Fig. 6b), supported by Pfams such as CobN (Fig. 6a), highlighting a distinct adaptation.

### GOTHAM DB: an Online Database for Exploration of Host-associated Proteins and Protein Domains

To make these analyses available to the community, we have developed an online database called **G**enes **o**f **T**raits of **H**ost-**A**ssociated **M**icrobes (GOTHAM database, https://ngdc.cncb.ac.cn/GeneAdaptionDB), which allows exploration of all enriched Pfam domains and AFCs identified in our various analyses.

Through the intuitive user interface, researchers can explore microbial adaptation patterns by filtering host associations or specific kingdoms, examining protein and domain enrichment at the genus and family level, and searching for specific Pfam domains or AFCs (Supplementary Fig. 22). The database provides detailed tabular data, including Pfam IDs/UniProt accessions, and the clade in which it was enriched.

GOTHAM database is a valuable resource for researchers who wish to explore and characterize novel functions associated with host interactions or gain insights into host adaptation strategies and mechanisms across specific taxonomic clades. For advanced computational analyses, GOTHAM provides free downloadable access to the full genome metadata and classification labels used in statistical tests.

## Discussion

This study aimed to systematically identify bacterial genes involved in adapting to multicellular host niches through extensive data analysis. Phylogeny-aware enrichment analyses uncovered thousands of putative host-associated protein clusters and domains across various genera and families, including well-characterized functions essential for host interactions, such as nitrogen fixation, nodulation, and virulence factors. Additionally, 70 domains with unknown functions and 80 uncharacterized AFC protein clusters were enriched across multiple genera, highlighting their high potential as candidates with important host adaptive roles. In contrast, no shared environmental functions were identified across diverse taxa. This may be due to the vast physical and chemical heterogeneity of non-host environments, making it difficult to detect patterns common to all. A comparative analysis of plant-associated and animal-associated bacteria further revealed numerous proteins and domains unique to each host kingdom, providing insights into host-specific adaptation mechanisms.

To validate our findings, we selected seven enriched genes without clear annotations and generated genome knockout mutants in multiple root-colonizing bacteria. Five out of seven mutant strains showed a significant reduction in rice root colonization, including both the rhizosphere and endosphere, compared to the wild type, while retaining their ability to grow in rich media. This suggests that our comprehensive approach successfully identified true host-associated proteins and protein domains. Follow-up experiments suggested that a new colonization factor (gen L) is involved in motility and in resistance to ROS; these are important functions that enable plant colonization.

### Pfam versus AFCs

We noted that AFCs had many more enriched “hits” as compared to Pfam analysis. This is partly due to the number of items in the database (24k entries in the Pfam database, versus 2.3 million non-singleton clusters in AFDB). Perhaps most importantly, the Pfam database covers domains and the AFC database covers whole proteins. This contrast between a portion of a protein, and a multi-domain protein considered holistically has effects on the outcome of enrichment tests. A domain, for example a DNA-binding domain, may or may not be part of a protein related to host-association– only the context of the other domains in the protein shape its holistic function. On the other hand, AFCs embody whole proteins (essentially an ordered permutation of domains) that are likely more specialized for a certain function like host-association, and therefore have a much clearer enrichment signal in host-associated genomes. Using both methods gives us a sense of whole proteins that are putatively involved in host-association, as well as a higher resolution view at the domain level.

### Role of Mobile Elements in Host-Associated Bacterial Adaptation

Most functions enriched in host-associated bacteria are not universally shared across clades adapted to different kingdoms. This is particularly evident when examining the top widespread Pfam domains and AFCs, which tend to be specific to genera with distinct host kingdom associations (Fig. 2a, 3a). For instance, genera such as *Paraburkholderia*, *Bradyrhizobium*, and *Mesorhizobium* exhibit numerous plant-associated functions, including nitrogen fixation and nodulation. Conversely, *Escherichia*, *Citrobacter*, and *Acinetobacter* are enriched in animal-associated functions like iron binding.

An interesting exception to this pattern is mobile elements, which are frequently found across bacteria adapted to both plant and animal hosts, spanning multiple distant taxonomic clades. For example, the DDE domain (PF13610), present in various transposases such as IS240, IS26, IS6100, and IS26^81^, is significantly enriched in gram-negative and -positive host-associated genera like *Bradyrhizobium*, *Burkholderia*, *Enterococcus*, *Escherichia*, and *Kocuria*. These mobile elements are often located near genes that may enhance bacterial fitness within host environments, as observed in the operons for Aerobactin siderophore, antibiotic and Mercury resistance (Fig. 4b,d-e). For example, *E. coli* that cannot use the Aerobactin siderophore (Fig. 4b) is highly defective in colonization of bladder and kidneys in a competition assay against wildtype *E. coli*^82^. It was also reported that antibiotic and Mercury resistance genes are often correlated and co-localized on the same genetic elements^83,84^.

The enrichment analysis applied in this study accounts for the phylogenetic structure of each taxonomic clade, favoring the identification of genes present in distant branches of the phylogenetic tree. This suggests that many of the identified genes were likely acquired through horizontal gene transfer, which plays a critical role in bacterial adaptation to host environments. Alongside host-adaptive genes, the mobile elements facilitating horizontal transfer were likely co-opted by recipient bacteria, enhancing their ability to thrive in host-associated niches. Previous studies have documented mobile elements carrying specific functions such as the Tn3 family, for their role in harboring antibiotic resistance genes^79^. Consistent with this, Tn3 was highly enriched in host-associated *Enterococcus*, *Escherichia*, and *Pseudomonas*. Our findings emphasize the crucial role of mobile elements in the adaptation of bacteria to host environments by facilitating the transfer of essential functions across diverse bacterial clades. On the other hand, we identified specific transposase and integrase domains that are depleted from hosts (Supplementary Figure 3).

### Comparison to previous GWAS with similar research questions

Previous studies have addressed similar questions. Pathfams^23^ identified pathogen-associated domain families, overlapping 34% with our enriched host-associated domains. Differences likely stem from differences in datasets and methodologies; Pathfams relied on the PATRIC database^42^, covering only 17% of our dataset, and used a hypergeometric test. In contrast, our study employed five statistical tests, including phylogeny-aware methods, for greater robustness.

Similarly, bacLIFE^24^ which focuses on prediction of lifestyle associated genes using three genera as case studies, while we analyzed 114 genera, offering broader coverage. A key distinction is the clustering method: bacLIFE used sequence-based clustering, while we used protein structure-based clustering for higher clustering sensitivity. Additionally, bacLIFE assigned lifestyles from species-level literature, whereas we used isolate-level genome metadata. bacLIFE also focused solely on gene presence/absence, while our approach, using methods like Evolink^27^, incorporated gene copy number, adding another layer of association detection.

A recent analysis revealed microbial gene modules that are linked to gut colonization^85^. The authors compared 9,475 representative mostly metagenome-associated genomes to compare gut microbes vs. environmental microbes. Using one statistical approach^86^ they defined 79 gut colonization factors and 23 multi-factor colonization modules. Two proteins were shown to be required for *E. coli* intestinal colonization in elegant experiments.

Overall, differences in methodologies, databases, and tools shaped the varied outcomes, with our approach integrating a comprehensive database, advanced statistical methods, and both structure based and domain-level analysis for robust conclusions.

### Limitations

While our systematic approach identified numerous Pfam domains and AFCs associated with host niches, several limitations warrant consideration. Genome labels were based on descriptions provided by sample collectors, which may be inaccurate or ambiguous, potentially leading to erroneous conclusions about domain or AFC enrichment. The vast quantities of data we collected hopefully drown out the occasional error. Additionally, the isolation site may not fully reflect a strain’s environmental adaptation. For example, a bacterium isolated from soil could still colonize a plant or animal but would be categorized as an environmental isolate in our analysis. To address these challenges, we employed five enrichment tests across three randomly sampled genomic datasets, and by validating our findings through computational and experimental methods.

Bioinformatic databases often lose relevance as time progresses. To keep GOTHAM updated and useful we plan to release future versions of the database that are based on the analysis of newly sequenced genomes, including from taxa that are not currently included, such as archaea, as well as newly developed analytical tools for enrichment detection.

### Summary

In conclusion, applying our computational approach to a massive bacterial genomic database allowed us to uncover numerous proteins and domains involved in the diverse mechanisms of bacterial host adaptation. These novel discoveries could pave the way for engineering beneficial microbes or developing new drugs targeting important host colonization factors, offering valuable insights for future research and applications in microbiology and biotechnology.

## Code Availability

All documented scripts are available in the GitHub repository at https://github.com/OfirSegev/Host-associated-project/.

## Acknowledgements

The project was mostly funded by the Israeli Science Foundation (Grants #1535/20, #3062/20) given to AL. In addition, it was partially supported the Israeli Ministry of Innovation, Science, and Technology (Grant #1001695377), and Israel Innovation Authority (Grant #81259), ICA in Israel, Israel Ministry of Agriculture (Grant 12-12-0008), the Volkswagen Stiftung (Grant ZN4041), and the Institute of Environmental Science of the Hebrew University of Jerusalem.

## Materials and Methods

### Retrieval of information on microbial genomes

A total of 931,343 genome sequences of bacterial isolates were obtained from 10 different public data sources^34–43^, including IMG, NCBI and PATRIC, updated to December 2021 (Supplementary Table 13). Genome sequences that contain information at the nucleotide level alone, were processed to generate the amino acid sequences using Prodigal^87^ version 2.6.3 (parameters: -a -f gff -c).

### Genome quality assurance

The genome sequences were filtered to include only high-quality genomes that met certain criteria. Specifically, genomes with completion below 95%, contamination above 5% using CheckM^88^ v1.1.3 (parameters: checkm lineage_wf -t 30 --tab_table), or N50 scores lower than 50,000 bp were removed. The remaining 397,116 genomes were used for further analysis. MAGs were also removed from the analysis as they may suffer from missing information and high contamination.

### Redundancy filtering

To minimize bias towards frequently sequenced species, such as well-known human pathogens, the genomes were filtered for redundancy. The nucleotide sequences were clustered based on sketch-based distance estimation using RabbitTClust^44^ (parameters: -k 21 - d 0.001). The genomes were clustered into 72,079 clusters. Each of these genome clusters was represented by a unique bacterial genome chosen by random, ensuring that no redundant or highly similar genomes were included in subsequent analyses.

### Genome labeling

Our dataset of genomes was characterized by highly heterogeneous metadata, often containing lengthy text strings that were impractical to parse manually. To classify each genome as host-associated or environmental, we manually labeled a subset of the most frequent metadata (“host” or “non-host”; “animal” or “plant”). Then, we implemented a natural language processing approach to encode the data, and trained a machine learning model to automatically classify metadata into a binary label. This model was used to classify the rest of the unlabeled genome metadata strings.

The raw metadata strings were labeled manually by a group of 10 degree-holding biologists who labeled the isolation metadata text (e.g. “human gut”) into one of “host”, “non-host”, and when information was available into “animal” or “plant”. The final training data set consisted of 23,354 manually labeled metadata descriptions: 18,663 labeled as host-associated and 4,691 as environmental. This dataset represented 70% of our genomes, as multiple genomes could share the same metadata description (e.g. a multitude of genomes had a host label of “*Homo sapiens*”, so one point of training data represents many genomes). We excluded metadata entries related to food, as food is processed and may not accurately represent host-niche colonization. Similarly, metadata entries from engineered environments (e.g., fermentors, wastewater remediation) were also removed from the dataset.

To encode the strings into proper input for machine learning, we one-hot encoded the raw metadata using TfidfVectorizer from scikit-learn^89^, with the parameter ngram_range=(1,2) . The output is a numerical matrix representing each metadata string, and each has an associated label. We then trained a logistic regression model to classify each input into a probability of being a binary label, using LogisticRegression from scikit-learn^89^, setting the hyperparameter C=5, with all other settings as standard. The model was trained with an 80-20 train-test split, achieving an accuracy and F1 score of 0.93. This trained model was then used to evaluate unlabeled metadata entries. To ensure the rigor of the labeling, we assigned those with prediction probability greater than 0.7 as either host-associated or environmental, discarding those near the classification threshold (probability of 0.5). Additional labels for host type (animal or plant) were assigned using the same approach.

### Division of genomes into taxonomic groups

To understand differences in gene content based on the environments in which the microbes live, rather than other taxonomic characteristics, the genomes were divided into taxonomic groups for separate analysis. We used GTDB-Tk^45^ (gtdbtk classify_wf, default parameters) to reveal the full taxonomy of each genome. Briefly, GTDB-Tk assigns taxonomy to microbial genomes by extracting 120 conserved marker genes, aligning them with HMMs, and placing genomes into a reference tree using maximum-likelihood methods. The resulting taxonomy annotation was then used to divide the genomes into separate genera and families.

### Protein domains clusters

Using the amino acid fasta files as inputs, protein domains were clustered and annotated using hmmscan^90^ (parameters: --domtblout) with e-value cutoff of 10e-5, minimum coverage of 70% and the Pfam hmm database as a reference^46^. These protein domain annotations were later used for the enrichment analysis at the protein domains level.

### Protein clustering to AFCs

All protein sequences were clustered at 80% coverage and 40% sequence identity using mmseqs (version 14.7e284), with the linclust^91^ subcommand (parameters: -c 0.8, -min-seq-id 0.4, –cov-mode 0). The representatives of the proteins were put into their own mmseqs database with the mmseqs subcommand createsubdb. In order to map the protein sequences to structures we considered predicting the protein structure de novo of the protein representatives, but this was untenable because of the scale and cost of GPU usage. To overcome this, we mapped the representative sequences to the Uniprot database^59^ (version 2022_3) using mmseqs search (parameters: –start-sens 1, –sens-steps 3, -s 7, –cov-mode 0, -c 0.85), using a highly conservative 85% coverage parameter. The output is a mapping from our protein representative database to Uniprot accession numbers. Furthermore, all of the protein structures in uniprot version 2022_3 are available and have been clustered structurally by using Foldseek^32^, and the pre-calculated clusters (90% structural overlap) are available^33^. This provided us with the structural clustering of all proteins in our genomes without computational overhead (Supplementary Table 4).

### Identification of host-associated proteins and protein domains

To identify proteins and protein domains that are associated with a host or lifestyle, we conducted five different enrichment tests: Scoary^25^ (parameters: -p=1.0), Pyseer^26^ (parameters: --lmm,--max-af=0.97), Evolink^27^ (presence absence, isolation forest), Evolink (counts, isolation forest) and Fisher exact test on each taxonomic clade. For the identification of enriched proteins, Alphafold clusters were used as input, while Pfam clusters were utilized for enrichment analysis on the protein domain level.

To minimize bias from random genome selection, the analysis was performed in three replicates, each using a randomly chosen representative genome from each genome cluster to form a non-redundant dataset. Enrichments consistent in at least two replicates were considered truly enriched.

Protein clusters or Pfams with prevalence of 97% or more among all genomes in a certain clade were removed from the Scoary analysis, due to lack of variance. A protein cluster/Pfam domain was considered significant by Scoary only if the q-value for Fisher’s exact test, which was calculated by the program was < 0.05, the ‘worst’ p-value from the pairwise comparison algorithm wasL< 0.05 and an odds ratio >1.5 or <0.67. For Pyseer’s Lmm, for each clade, we generated a q-value using fdrcorrection from statsmodels.stats.multitest for both chi-square and likelihood ratio test p-values. A Protein/domain was considered significant by Lmm if the chi-square q-value was < 0.01 as well as the likelihood ratio test q-value and an odds ratio >1.05 or <0.95. The Fisher test was performed using Scoary’s output. The thresholds for a significant hit were a bit stricter, where q-value <0.001 and odds ratio >2 or <0.5. For Evolink tests, the threshold for identifying gene families with significant associations is determined by the upper bound of maximal differences of the sorted outlier scores and was determined automatically by the program itself.

Scoary, Pyseer, Evolink (presence/absence version) and Fisher consider the presence or absence of a protein/domain, where Evolink (counts version) considers gene copy-number data. In addition, for each clade, a phylogenetic tree consisting of all clade members was constructed using PhyloPhlAn (version 3.0.60)^92^. The parameters were d: phylophlan, min_num_entries: was set at 90% of total genus size, diversity: medium, submat: pfasum60, trim: greedy, Remove_fragmentary_entries, fragmentary_threshold: 0.5, subsample: twentyfivepercent, scoring_function: trident, not_variant_threshold: 0.9, gap_perc_threshold: 0.7. The f flag took in a conFig. file describing the programs and parameters from the following pipeline: diamond (makedb), diamond (blastx --quiet --threads 1 --outfmt 6 --more-sensitive --id 50 --max-hsps 35 - k 0), diamond (blastp --quiet --threads 1 --outfmt 6 --id 50 --max-hsps 35 -k 0 --fast), mafft (-- memsavetree --anysymbol --thread -1 --auto --retree 1 --quiet), trimal (-gappyout), and finally FastTreeMP (-quiet -pseudo -spr 4 -mlacc 2 -slownni -fastest -no2nd -mlnni 4 -lg)

This tree was later used as an input for all tests, as they take into account the phylogenetic distance between the genomes, which is calculated based on the clade’s tree structure. Each clade contained closely related unique genomes that were randomly sampled from the corresponding genome cluster. We repeated each test three times, randomly sampling different genomes for each repetition. AFC protein clusters/Pfams were considered to be significant in a certain test if at least two samples identified the same protein clusters/Pfams that passed the assigned thresholds.

The analysis was performed solely on clades with sufficient number of genomes and high label entropy: H = −(p1*log_2_p1 + p2*log_2_p2). Entropy serves as a metric of randomness, reflecting the diversity of labels within a specific clade. In this context, if a clade predominantly comprises genomes of a certain label, it would display low entropy, as a result, there would be insufficient representation of other labels for meaningful comparison. Clades with over 100 genomes were subjected to a minimum entropy threshold of 0.25, those with a genome count ranging from 50 to 100 adhered to a minimum entropy of 0.5, and for clades with fewer than 50 genomes, a minimum entropy threshold of 0.8 was applied. Clades comprising less than 20 genomes were automatically excluded from the analysis.

### Advantages and disadvantages of the different algorithms

In general, Fisher Exact Test is the most promiscuous test (Supplementary Figure 1b) and will identify correlations between a pfam/protein and host-association independently of any phylogenetic information. Namely, it does not correct for phylogenetic relatedness or population structure. Therefore, it may identify false positive associations due to enrichment of a certain pfam/protein in one branch of the gene tree that happens to have the same host-association label, independently of real adaptation to the niche (e.g. due to short evolutionary time). Scoary scores the components of the pan-genome for associations to observed phenotypic traits while accounting for population structure. It supports only binary data (presence/absence) and not copy number variations as other tools do. Scoary implements the pairwise comparisons algorithm to control spurious associations from stratified populations, which is effective for controlling for bias from clonal sampling of bacterial genomes. It was tested numerous times by other groups and is the most cited approach among the ones we used, except for the classic Fisher exact test. Pyseer uses linear models with fixed (generalized linear models) or mixed effects (linear mixed models) to estimate associations, accounting for strong confounding due to population structure—an essential feature for clonal microbial organisms. The complex models it uses can be computationally intensive and require substantial memory and processing power. The effectiveness of population structure correction relies on the accuracy of phylogenetic reconstructions. We observed the lowest number of predictions for Pyseer (Supplementary Figure 1b) showing potentially reduced sensitivity for the current GWAS application. Evolink is a relatively new method that also corrects for phylogenetic structure and provides an index that quantifies the strength of gene-pheotype association. It is one of the fastest methods in terms of computation time. Like other methods it is highly dependent on the accuracy of the phylogenetic tree. Evolink developers argue that their method had higher precision than Pyseer on empirical dataset^27^. In addition, this is the only tool which allows us to also use copy number data which can be important for quantitative phenotypes.

To elucidate the overall trends and attributes of enriched Pfam domains across distinct groups, we conducted a gene ontology (GO) analysis. First, each Pfam domain matched to its corresponding InterPro ID sourced from the InterPro database^81^. Each InterPro ID is associated with a specific set of GO terms, thus allowing for the correlation between Pfam domains and their respective GO term annotations^93,94^.

It is noteworthy that while the majority of Pfam domains were successfully linked to an InterPro ID, 1% of the domains remained unmatched. Additionally, 73% of the InterPro IDs that are linked to a Pfam domain lacked any GO term annotations, resulting in a significant proportion of Pfam domains remaining unannotated with GO terms.

To identify the most representative GO terms for each distinct habitat within each clade, we used the survival function of the hypergeometric distribution for each clade independently to generate corresponding p-values (scipy.stats.hypergeom.sf(k-1, M, n, N)^95^). The parameters for each GO term were defined as follows: k is the number of occurrences of the specific GO term within the enriched results, N is the total number of GO terms in the root GO category that were mapped to the enriched results, n is the number of occurrences of the GO term in the overall population of the relevant clade, and M is the total number of GO terms in the root GO category that were mapped to all Pfam domains encoded by the entire population of the clade.

In this analysis, only Pfam domains found to be enriched in two or more different enrichment tests within a particular clade were considered enriched. The resulting p-value reflects the probability that a randomly sampled GO term would equal or exceed the observed value derived from the enriched Pfam domains within that clade. To control for multiple hypothesis testing and manage the false discovery rate, we applied FDR correction to the p-values (smm.fdrcorrection with alpha=0.05). Additionally, fold change was calculated for each GO term that met the threshold of a q-value less than 0.05. The fold change was calculated as the ratio of the proportion of a specific GO term in the enriched sample set to its proportion in the overall population, expressed as (k/N)/(n/M). In each set, the proportion represents the number of Pfam domains in a specific category divided by the total number of domains within the corresponding root category. The package ggplot2 was used for visualization^96^.

### Gene ontology analysis of Alphafold protein clusters

Since the AFCs have Uniprot accessions, the associated GO annotations are readily available^59^. Each AFC representative was considered labeled with a GO term only if at least 75% of the structural cluster’s members had the same GO term associated with it. Out of all enriched results, 56% were mapped to a GO using this threshold. The rest of the GO analysis was performed using the hypergeometric distribution by applying the exact same methodology as discussed at the gene ontology analysis of Pfam domains.

### Operon clustering

Enriched Pfams from .gff files were queried, and scaffolds were segmented into 20,000 bp chunks. Pfam co-occurrence within these chunks was used to calculate pairwise correlations across all host-associated genomes in a given genus. Pfam pairs with correlation values above 0.1 were clustered, allowing for redundancy of Pfam domains in multiple clusters. Singleton clusters were excluded. The clusters were further assessed for the mean distance between the end coordinates of each gene, quantifying proximity. Additionally, cluster directionality was evaluated by determining the percentage of genes oriented in the same direction. Clusters with a mean distance under 3000 bp and directionality greater than 65% were classified as operons. For the operons presented, all genes within each operon were mapped to AFCs to assess their enrichment within the overall enrichment results.

### Principal coordinates analysis

To systematically and quantitatively validate these results, we performed dimensionality reduction using Principal Coordinates Analysis (PCoA) on the genomes in each analyzed genus and family. This analysis utilized distance matrices generated from the counts of AlphaFold Clusters (AFCs) or Pfams in each genome. We then repeated this analysis using only the subset of enriched AFCs or Pfams, rather than the complete set of all AFCs and Pfams in each clade. Our hypothesis was that this latter PCoA would achieve better separation between “host-associated” and “environmental” or “animal-associated” and “plant-associated” labels. We evaluated the results using a logistic regression model by measuring the change in F1 score and the change in Akaike Information Criterion (AIC) between models based on the full set of AFCs/Pfams and those based on the enriched subset. F1 score is defined as 2*(TP) / [2TP+ FP + FN], where TP, FP, and FN are true positives, false positives, false negatives, respectively. AIC is an estimator of prediction error, defined as AIC = 2k - 2ln(L), where k is the number of estimated parameters in the model, and L is the maximized value of the likelihood function for the model.

Examples of PCoA results for representative subsets are provided in the supplementary figures (Supplementary figures 13-16).

In the comparison between host-associated and environmental bacteria, we found that using the enriched Pfams resulted in a reduction in AIC without a decline in F1 score in 110 out of 114 genera (96.5%) and 62 out of 66 families (93.9%) (Supplementary Table 8). A similar trend was observed with AFCs, with 105 out of 114 genera (92.1%) and 63 out of 66 families (95.4%) showing a reduction in AIC. These findings suggest that the enriched proteins and domains are significant factors in defining host association within these clades.

In the comparison between animal-associated and plant-associated bacteria, most clades also showed improvement when PCoA was applied using only enriched genes. Specifically, 27 out of 28 genera (96.4%) and 31 out of 35 families (88.6%) showed improvement when using Pfam domains. A similar trend was observed with AFCs, with 26 out of 28 genera (92.9%) and 32 out of 35 families (91.4%) demonstrating improvement.

### Candidates selection for experimental validation

Six Pfam domains that correspond to six AFCs, identified as enriched in host-associated bacteria, were selected for experimental validation. The selection criteria for these Pfam domains and AFCs were based on several factors, including their presence in at least one of the bacterial strains maintained by our collaborators (Vittorio Venturi’s lab), the availability of a genetic manipulation system for the bacteria, limited annotation information, association with bacteria known to interact with plants, and a low copy number of the gene in the genome allowing straightforward gene knockout performance. Subsequently, knockout mutations were induced in seven genes housing the candidate Pfam domains and that were clustered with enriched AFC (Supplementary Table 9). Following this, several experiments were conducted to assess the impact of the mutations on bacterial behavior.

### Bacterial strains, growth conditions and growth curves generation

The bacterial strains used in this study were as follows:

*Kosakonia oryzae* KO348^97^

*Pseudomonas fuscovaginae* UPB0736^98^

*Dickeya zeae* DZ2Q^99^

*Pseudomonas capeferrum* WCS358^100^

All strains were grown in liquid Luria-Bertani [LB-10 g tryptone, 5 g yeast extract, and 10 g NaCl in 1 liter], Tryptic Soy Broth [TSB-30 g of commercially available Tryptic Soy Broth powder in 1 liter] or M9 medium [6.0 g NaLHPOL, 3.0 g KHLPOL, 0.5 g NaCl, and 1.0 g NHLCl, 1 mL of 1 M MgSOL, 0.1 mL of 1 M CaClL, and 2 g/L glucose in 1 liter] at 30°C under moderate shaking (200 rpm). All media were sterilized by autoclaving at 121°C for 15–20 minutes and stored at appropriate conditions until use. For solid media, 15 g/L of agar was added. When required, antibiotics for tested strains growth were added at the following concentrations: nitrofurantoin (Nf) 100 µg ml^−1^ for *Pseudomonas capeferrum* WCS358 and *Pseudomons fuscovaginae* UPB0736, and rifampicin (Rif) 100 µg ml^−1^ for *Kosakonia oryzae* KO348 and *Dickeya zeae* DZ2Q.The mutants of each strain (carrying a knock-out mutation) have been grown using 100 µg ml^−1^ kanamycin (Km) as antibiotics. *E. coli* DH5α was routinely grown at 37 °C in LB broth and antibiotics were added when required at the following concentrations: gentamicin (Gm) 10 µg ml^−1^, tetracycline (Tc) 10 µg ml^−1^.

To evaluate the growth dynamics of the tested strains, growth curves were performed. Subcultures of specified strains were grown overnight at 30°C, diluted in fresh medium to an OD600 of 0.1, inoculated into conical flasks, and grown at 30°C with 200 rpm. All the strains were cultured either in TSB or M9 supplemented with glucose 0.2%. Growth was monitored by measuring optical density at 600 nm (OD600) at regular intervals using a spectrophotometer over 24 hours. Data were collected in five replicates to ensure reproducibility (Supplementary Table 12).

### Mutant construction

Deletions of each target gene (geneA, geneB, geneD, geneF, geneG, geneI, geneL) were generated using the pEX19Gm plasmid as described previously^101^. Briefly, each gene sequence, synthesized by Twist bioscience company (South San Francisco), is listed in the Supplementary Dataset 1. The design of the constructs was performed as follows: internal fragments of 200 bp from each gene of interest were deleted and replaced with a restriction site (BamHI, SalI or PstI) in order to clone inside the kanamycin (Km) gene cassette previously extracted from pUC4K vector (Addgene, Watertown, MA). Sequentially the fragments were excised with the following restriction enzyme couples: EcoRI and HindIII or KpnI and HindIII or BamHI and HindIII, and cloned in the corresponding site in pEX19Gm. The resulting pEX19Gm-derivative plasmids were introduced by biparental conjugation in the corresponding host genomes. This method involves homologous recombination to introduce mutations and sucrose counterselection to eliminate cells that have not undergone double-crossover events, ensuring the selection of mutants with precise gene deletions. Clones with a chromosomal insertion of the pEX19Gm plasmids were selected on LB agar plates supplemented with 40 µg ml^−1^ Gm, 100 µg ml^−1^ Km and 100 µg ml^−1^ Rif or Nf. Plasmid excision from the chromosome was subsequently selected on LB agar plates supplemented with 15% (w/v) sucrose. All the mutants were verified by PCR using primers (Supplementary Table 14) specific to the genomic DNA sequences upstream and downstream from the targeted genes.

For the deletion of genes in *Kosakonia oryzae* KO348, a technical approach based on the plasmid incompatibility principle was employed^102,103^. In summary, the synthesized gene cassettes were cloned into the pLAFR3-TcR vector and introduced into *Kosakonia oryzae* KO348 WT via conjugation. Subsequently, a second plasmid, pPHJ1J-GmR (Addgene, Watertown, MA) was conjugated into the *Kosakonia* (pLAFR mutants), and transconjugants were selected on plates containing 40 µg/ml Gm and 100 µg/ml Km. PCR analysis was conducted to verify all transconjugants.

For complementation analysis of the *Pseudomonas capeferrum* WCS358 I and L, the full-length coding region of each gene was amplified using primers listed in Supplementary Table 14. The PCR products were digested with restriction enzymes and cloned into the expression vector pLAFR3^104^, which had been digested with the same enzymes. The resulting complementation constructs were introduced into the corresponding mutants via bi-parental mating, and transformants were selected based on Km^R^ and Tc^R^ resistance. PCR analysis confirmed the successful integration.

### Plant colonization experiments

Baldo rice seeds were surface sterilized with 10% bleach for 10 minutes, followed by five washes with sterile water. The seeds were pre-germinated for 8 days in a dark room at 30°C, then grown for 4 days in a greenhouse under a 25°C temperature and a 16-hour photoperiod to ensure uniform growth.

All the bacterial strains (WT and relative knock-out mutants) used in this study were grown on TSB to an optical density at 600 nm (OD600) of 0.8 and 12-day-old germinated Baldo rice plantlets were then submerged in this bacterial suspension for 1 h and transferred independently to a tube containing Hoagland’s semi solid solution ^105^.

Inoculated plantlets were watered and grown for 6 days in a greenhouse under controlled conditions (25°C, 16-hour photoperiod).

#### Rhizosphere Microbial Recovery

Roots were carefully removed from the growth medium to minimize damage. Excess Hoagland’s semi solid medium was gently shaken off. Roots with adhered rhizosphere soil were placed in sterile 50 mL conical tubes containing phosphate-buffered saline (PBS) and vortexed at high speed for 5 minutes. The samples were then sonicated in an ultrasonic water bath (40 kHz, 100 W) for 10 minutes. Roots were removed for endosphere recovery, and the remaining solution was centrifuged at 3750 × g for 10 minutes to pellet rhizosphere particles. The supernatant was discarded, and the pellet was resuspended in 5 mL of PBS. Serial dilutions of this suspension were plated on selective media to calculate CFU/g.

#### Endosphere Microbial Recovery

Roots were surface sterilized by sequential treatment with 50 mL ethanol (75%) for 2 minutes, sterile water, 50 mL bleach (50%) for 2 minutes, sterile water, and another 50 mL ethanol (75%) for 2 minutes. This was followed by six washes with sterile water. Sterility controls were performed as previously described by Bertani et al^97^. The sterilized roots were macerated in 2 mL of PBS, and serial dilutions of the macerate were plated on TSA containing selective antibiotics. Plates were incubated at 30°C for 24 hours, and CFU/g was calculated.

For the selective recovery of mutant strains, kanamycin (100 μg ml^-1^) was used. For *Kosakonia oryzae* and *Dickeya zeae*, rifampicin (100 μg ml^-1^) was utilized, while *Pseudomonas capeferrum WCS358* and *Pseudomonas fuscovaginae* were recovered using nitrofurantoin (100 μg ml^-1^).

### Swimming assay

*Pseudomonas capeferrum* WT, gene mutants, and complemented strains overnight cultures were grown as described above and diluted to a final density of 1 × 109 CFU/ml. After carefully placing 3 μl suspension droplets in the centre of soft swimming plates (M9 medium, 0.3% agar), plates were incubated at 30°C and evaluated after either 1 or 2 days by measuring the diameter of the swimming zone (Supplementary Table 9).

### H2O2-kill assay

H2O2-kill assay of *Pseudomonas capeferrum* WCS358 WT vs mutants WCS358 ΔI, WCS358 ΔL, complemented WCS358 ΔI(pBBR::I) and WCS358 ΔL(pBBR::L) under 2mM H2O2 was performed. Overnight cultures in TSB were inoculated into ME medium [20 g/L glucose, 12 g/L sodium citrate, 7 g/L (NH4)2SO4, 0.50 g/L K2HPO4·3H2O, 0.50 g/L MgSO4·7H2O, 0.04 g/L FeCl3·6H2O (0.15 mM Fe3+), 0.01 g/L MnSO4·H2O, and 0.15 g/L CaCl2, pH 7.5] and grown at 30°C for 12 h. Cells were then treated with 2 mM H2O2, and after 15 min of incubation, samples were diluted and plated on Tryptic soy broth agar (TSA), and incubated at 30°C for 16-18 h. Serial dilutions of recovered cells were plated, and cell survival was determined by counting CFU and is shown as the mean value ± standard error from five independent experiments (Supplementary Table 9).

