## Supplementary Material and Figures for "The Genetic Basis of Bacterial Adaptation to Hosts"

### Supplementary Information

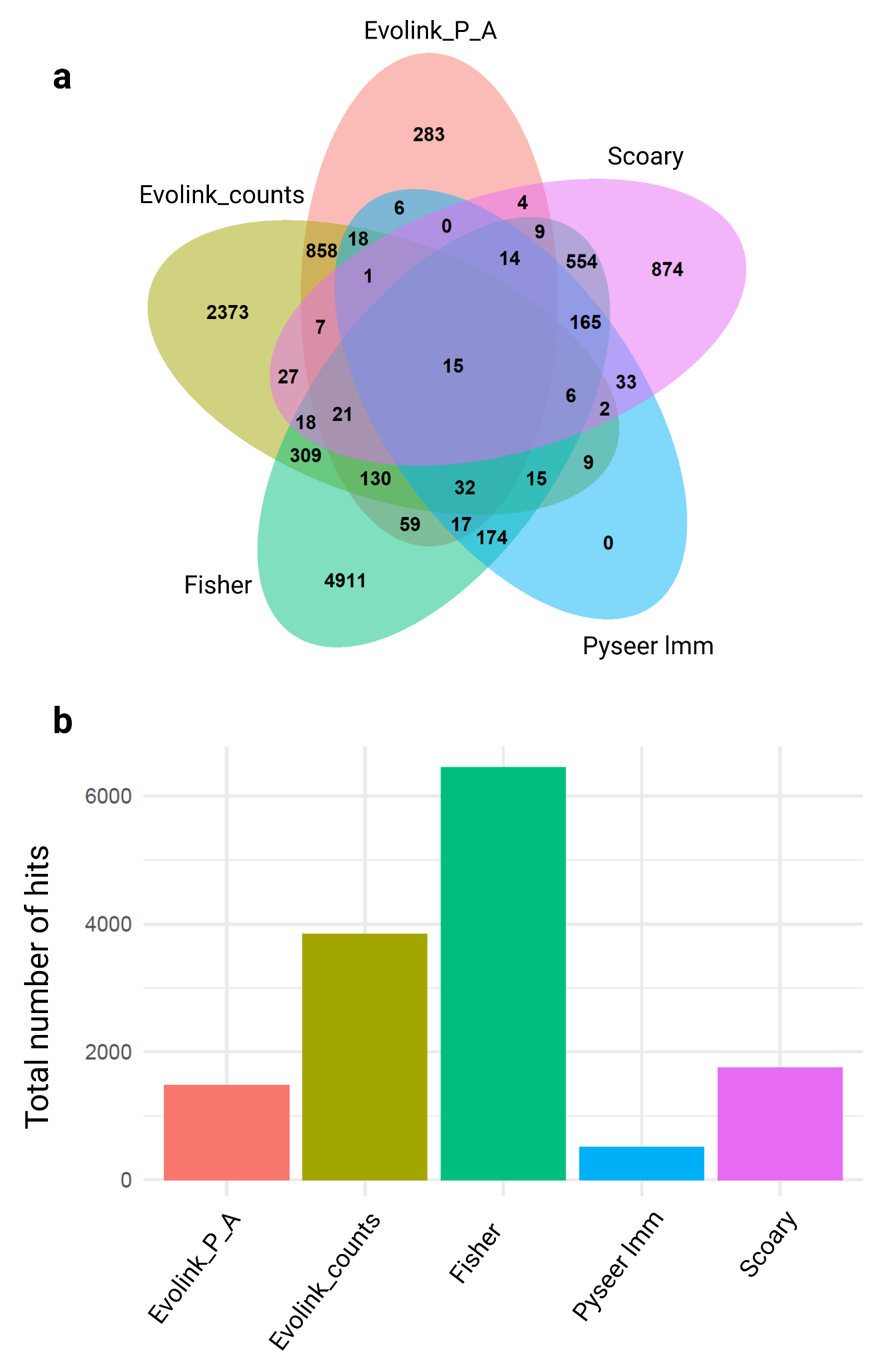

**Supplementary Figure 1: Pfam Enrichment Results by Five Distinct Enrichment Tests.**

**a, Overlap of enriched Pfams across different tests.** A Venn diagram illustrates the counts of host-associated Pfams identified by each enrichment test when taxonomic clades were grouped at the genus level. **b, The number of enriched Pfams identified by each enrichment test.** The x-axis represents the five different enrichment tests employed, while the y-axis indicates the number of host-associated Pfams enriched at the genus level.

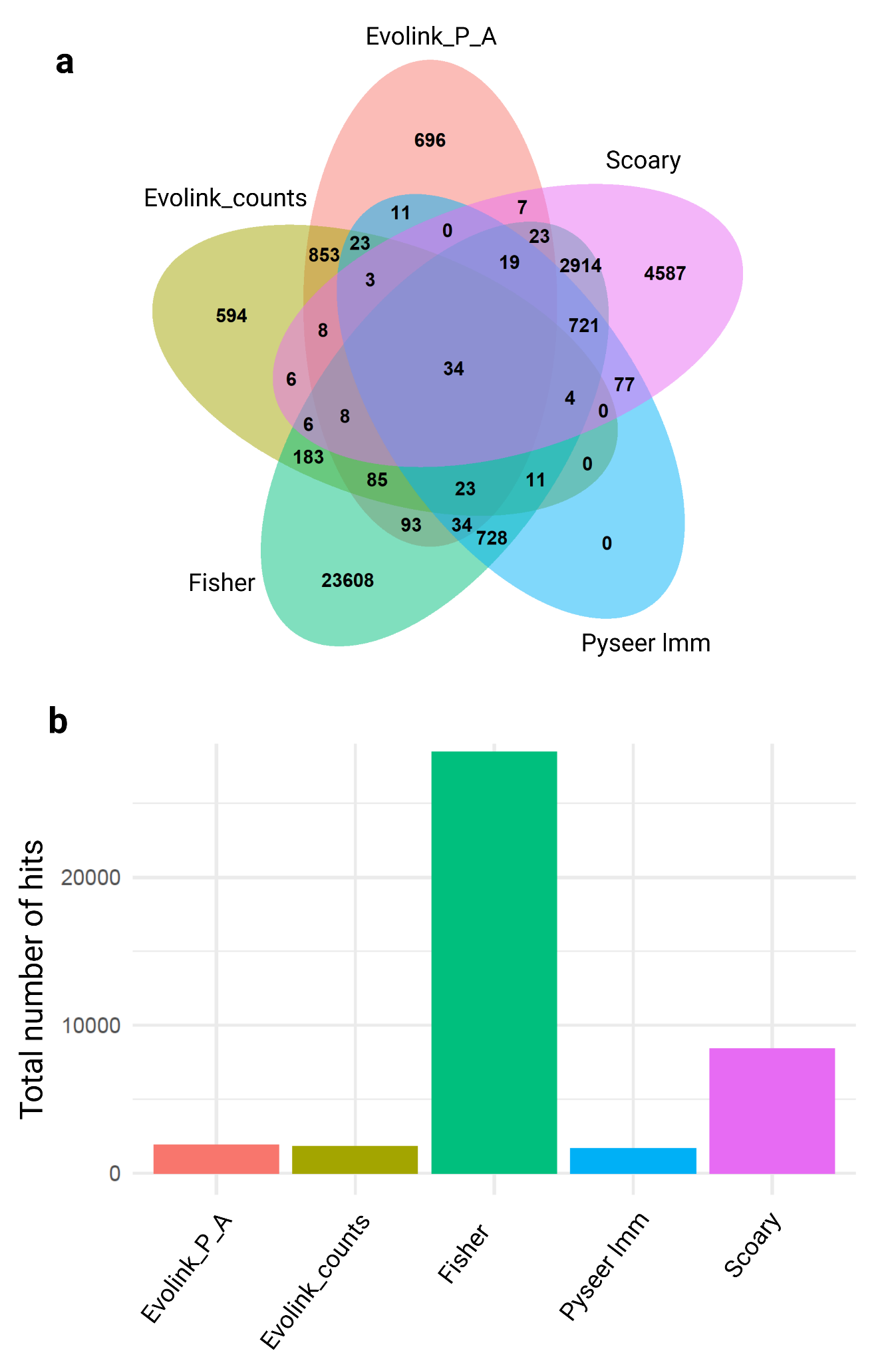

**Supplementary Figure 2: AFC Enrichment Results by Five Distinct Enrichment Tests.**

**a, Overlap of enriched AFCs across different tests.** A Venn diagram illustrates the counts of host-associated AFCs identified by each enrichment test when taxonomic clades were grouped at the genus level.

**b, The number of enriched AFCs identified by each enrichment test.** The x-axis represents the five different enrichment tests employed, while the y-axis indicates the number of host-associated AFCs enriched at the genus level.

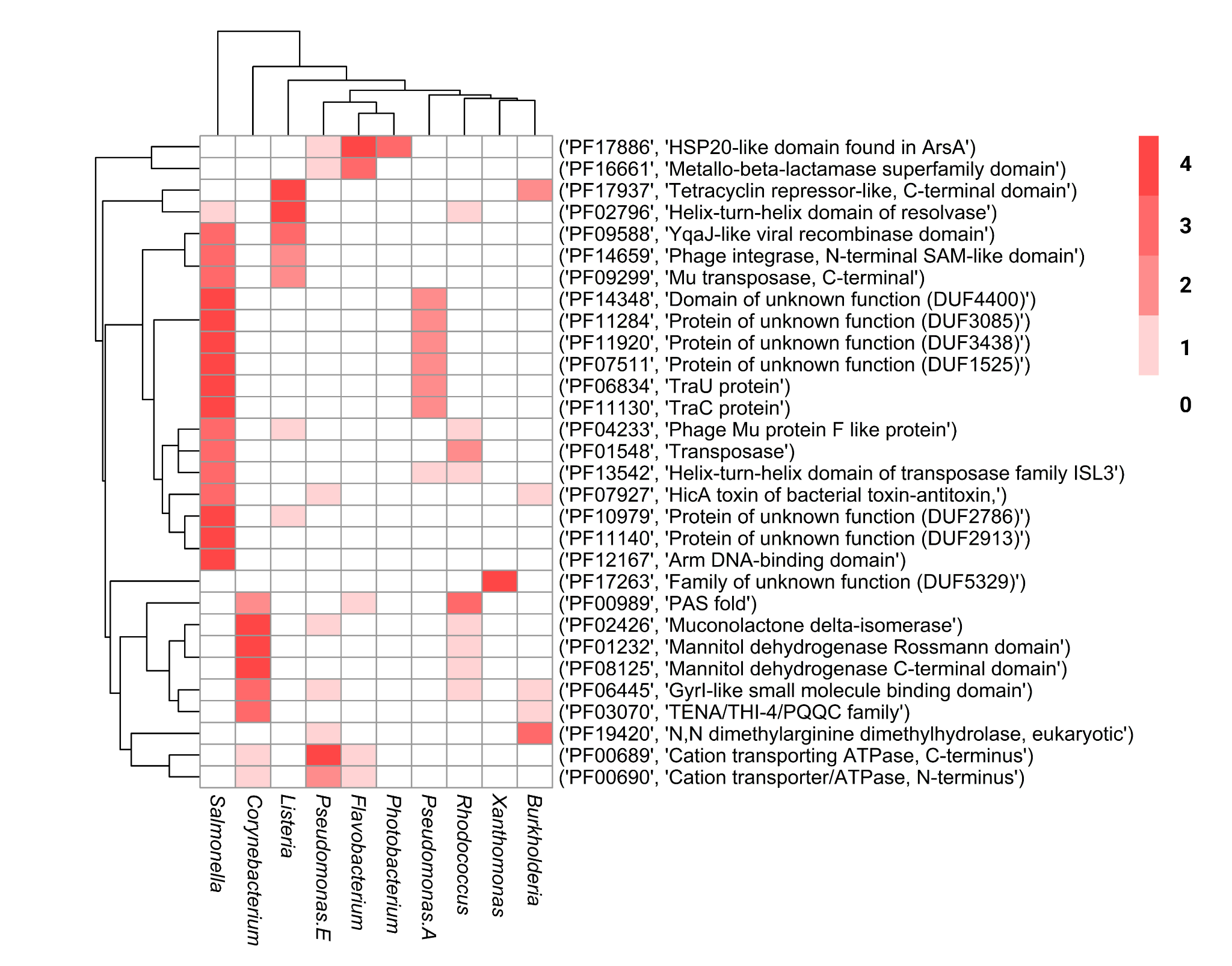

**Supplementary Figure 3: Heatmap showing the depletion of host-associated Pfam domains across 10 bacterial genera.** Each row represents a Pfam domain (Pfam ID and description), and columns correspond to genera. Red shading indicates the number of tests identifying these domains as significantly enriched in environmental bacteria within a clade, with white indicating no enrichment.

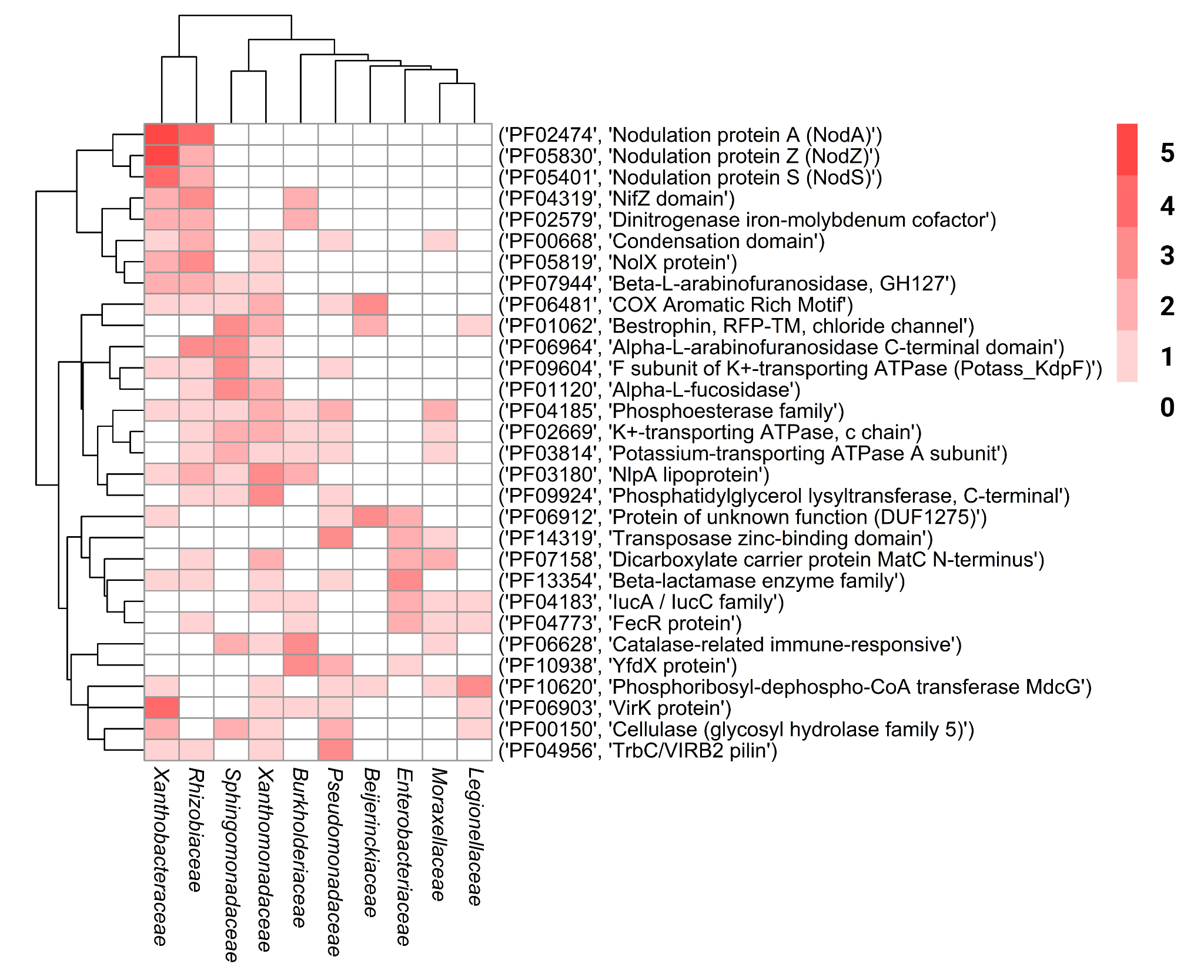

**Supplementary Figure 4: Heatmap showing the enrichment of host-associated Pfam domains across 10 bacterial families.** Each row represents a Pfam domain (Pfam ID and description), and columns correspond to families. Red shading indicates the number of tests identifying these domains as significantly enriched in host-associated bacteria within a clade, with white indicating no enrichment.

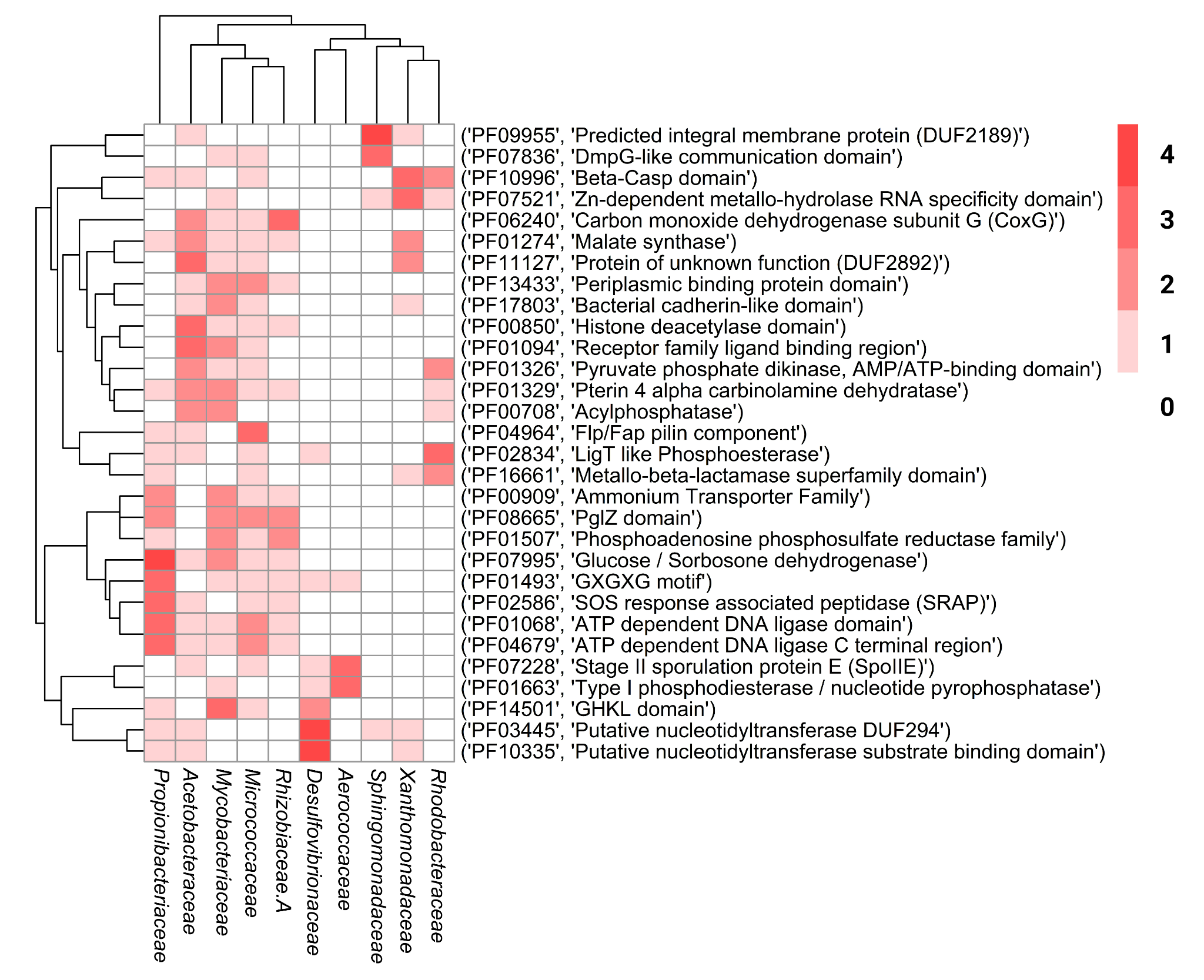

**Supplementary Figure 5: Heatmap showing the depletion of host-associated Pfam domains across 10 bacterial families.** Each row represents a Pfam domain (Pfam ID and description), and columns correspond to families. Red shading indicates the number of tests identifying these domains as significantly enriched in environmental bacteria within a clade, with white indicating no enrichment.

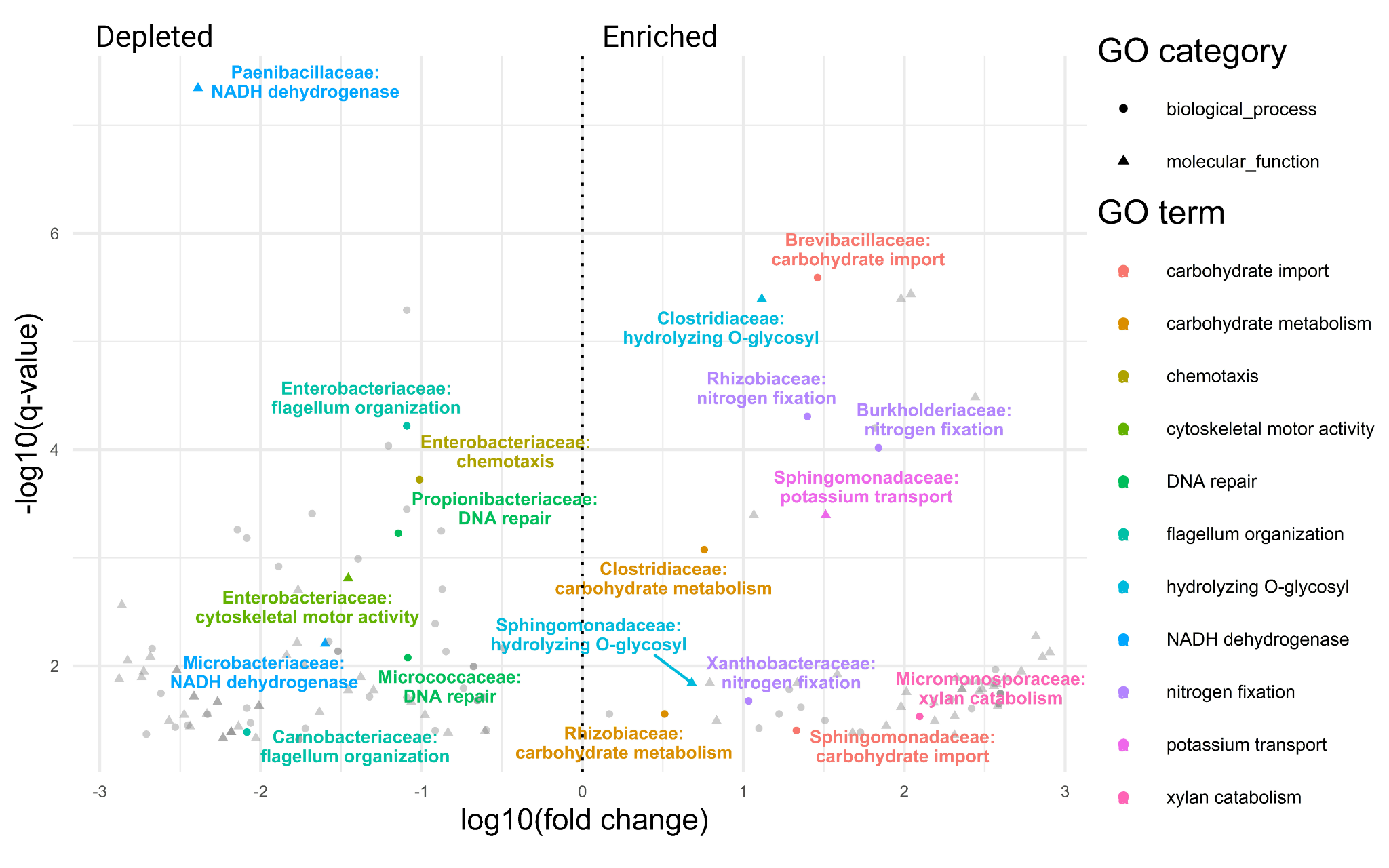

**Supplementary Figure 6: Volcano plot highlighting highly enriched host-associated functions based on Pfam domains at the family level.** The y-axis represents the significance of enrichment (-log10 FDR-corrected q-value), while the x-axis shows the fold change (log10 fold change) for each mapped GO term. Colors highlight specific key functions. Functions located to the right of the dotted line are enriched in host-associated bacteria, while those on the left are depleted. To improve readability, "carbohydrate metabolic process" was abbreviated to "carbohydrate metabolism", "xylan catabolic process" to "xylan catabolism", "hydrolase activity, hydrolyzing O-glycosyl compounds" to "hydrolyzing O-glycosyl", "NADH dehydrogenase (ubiquinone) activity" to "NADH dehydrogenase", “bacterial-type flagellum organization" to "flagellum organization", "P-type potassium transmembrane transporter activity" to "potassium transport" and “phosphoenolpyruvate-dependent sugar phosphotransferase system” to “carbohydrate import”.

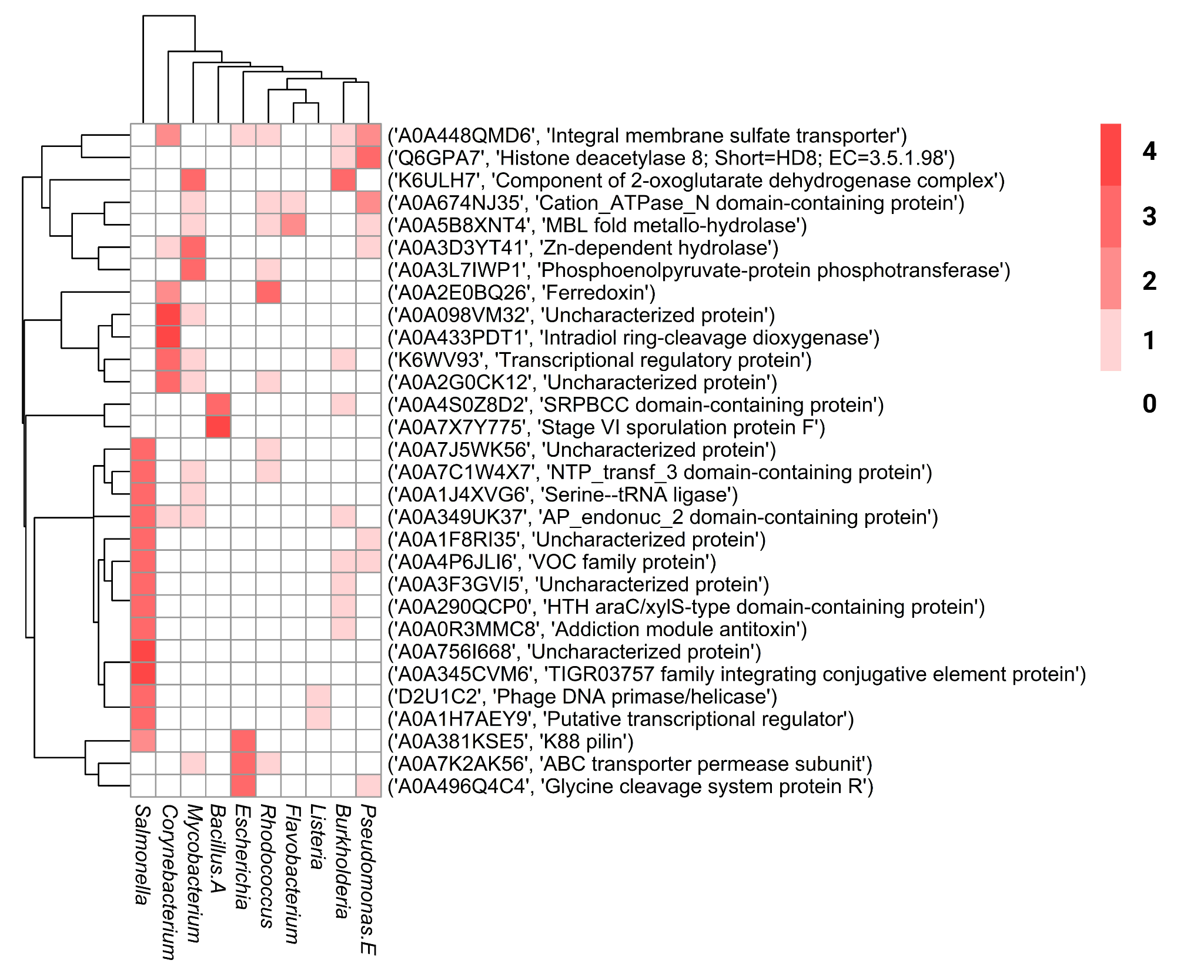

**Supplementary Figure 7: Heatmap showing the depletion of host-associated AFCs across ten bacterial genera.** Each row represents an AlphaFold cluster (UniProt accession and description), and columns correspond to genera. Red shading indicates the number of tests identifying these proteins as significantly enriched in environmental bacteria within a clade, with white indicating no enrichment.

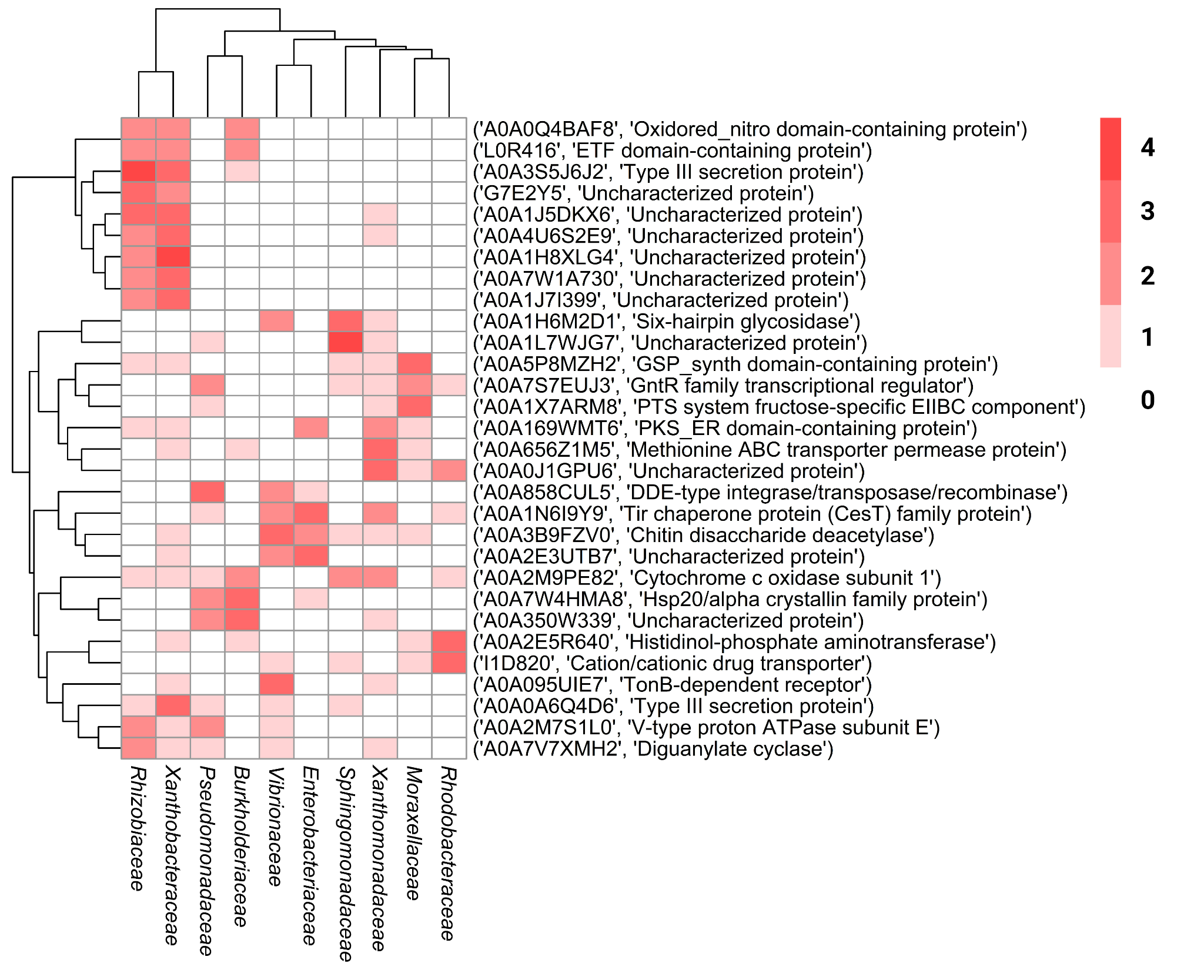

**Supplementary Figure 8: Heatmap showing the enrichment of host-associated AFCs across 10 bacterial families.** Each row represents an AlphaFold cluster (UniProt accession and description), and columns correspond to families. Red shading indicates the number of tests identifying these proteins as significantly enriched in host-associated bacteria within a clade, with white indicating no enrichment.

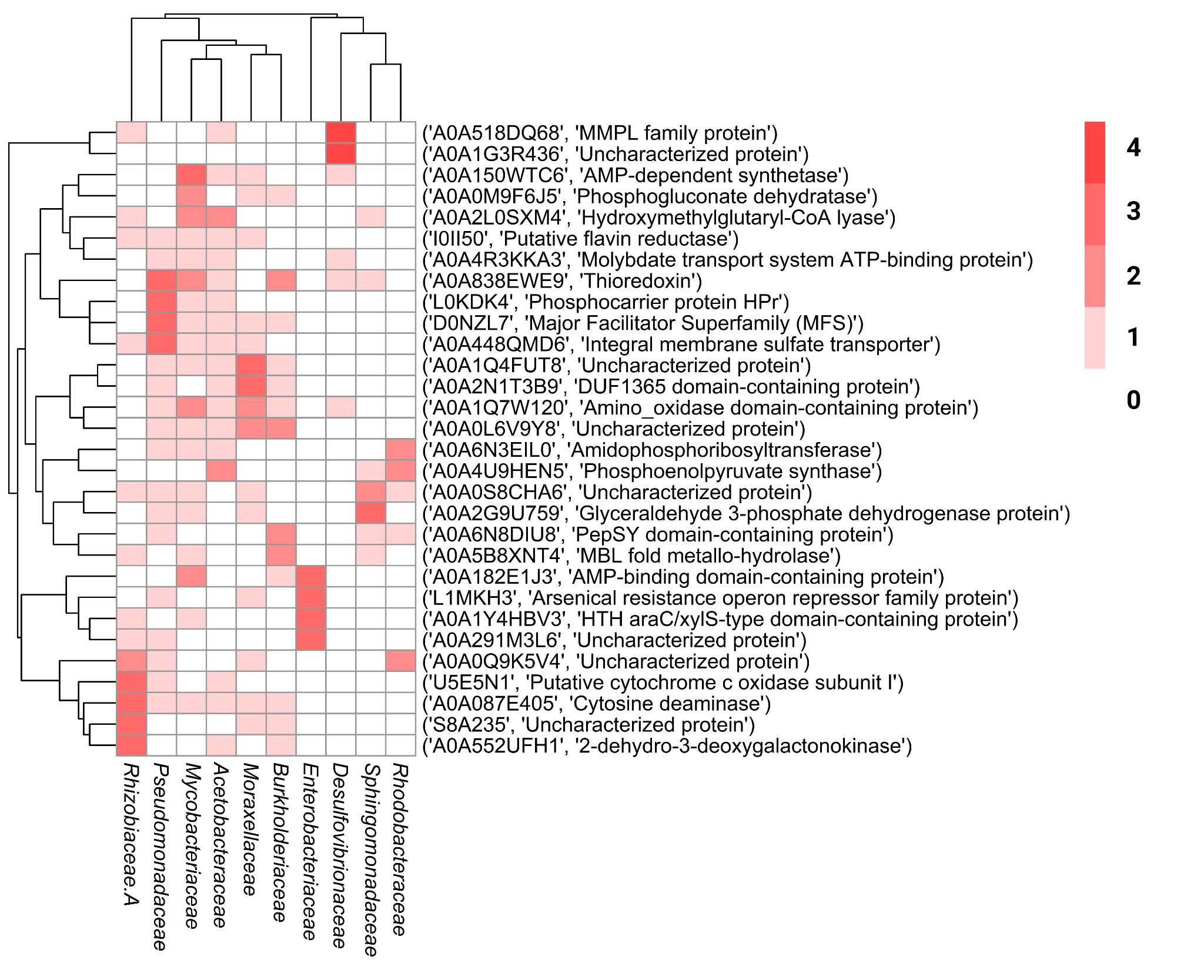

**Supplementary Figure 9: Heatmap showing the depletion of host-associated AFCs across 10 bacterial families.** Each row represents an AlphaFold cluster (UniProt accession and description), and columns correspond to families. Red shading indicates the number of tests identifying these proteins as significantly enriched in environmental bacteria within a clade, with white indicating no enrichment.

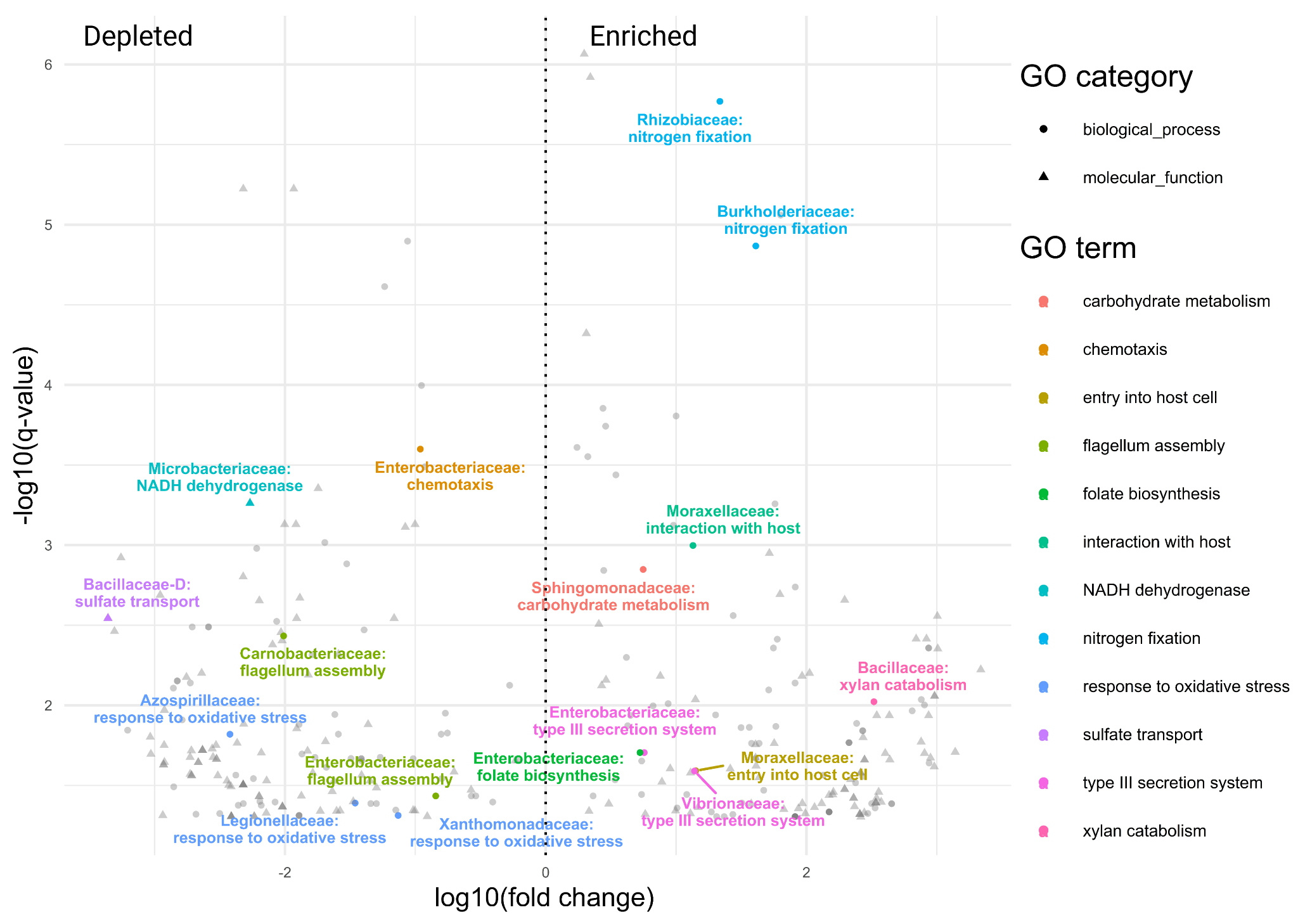

**Supplementary Figure 10: Volcano plot highlighting highly enriched host-associated functions based on AFCs at the family level.** The y-axis represents the significance of enrichment (-log10 FDR-corrected q-value), while the x-axis shows the fold change (log10 fold change) for each mapped GO term. Colors highlight specific key functions. Functions located to the right of the dotted line are enriched in host-associated bacteria, while those on the left are depleted. To improve readability, "protein secretion by the type III secretion system" was abbreviated to "type III secretion system", "carbohydrate metabolic process" to "carbohydrate metabolism", "biological process involved in interaction with host" to "interaction with host", ”symbiont entry into host cell” to “entry into host cell”, "NADH dehydrogenase (ubiquinone) activity" to "NADH dehydrogenase", “folic acid-containing compound biosynthetic process" to "folate biosynthesis", “secondary active sulfate transmembrane transporter activity” to "sulfate transport" and "bacterial-type flagellum assembly" to "flagellum-dependent motility".

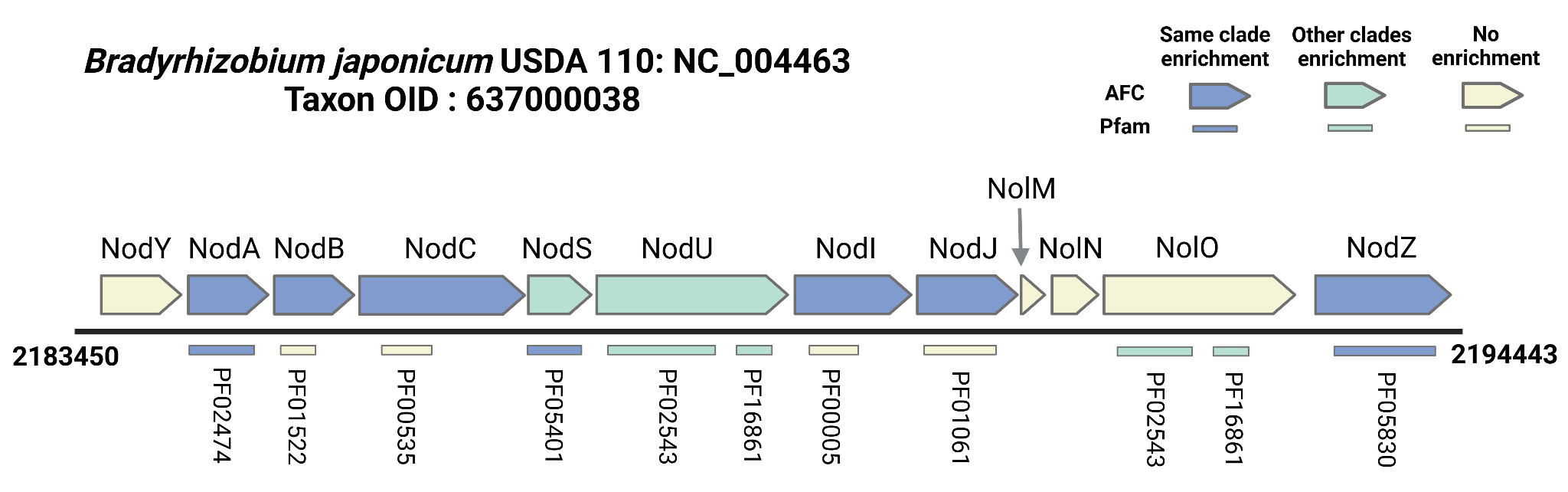

**Supplementary Figure 11: Illustration of nodulation operon found in *Bradyrhizobium japonicum* USDA 110.** Each arrow represents a gene located in the genome, with the bold numbers on the sides, indicating the genomic location within the contig. Genes highlighted in blue indicate enrichment within the same genus as the depicted strain (*Bradyrhizobium*), while teal indicates enrichment in other genera only. AFC enrichment is shown by the colored genes, and Pfam enrichment is indicated by the lines adjacent to those genes.

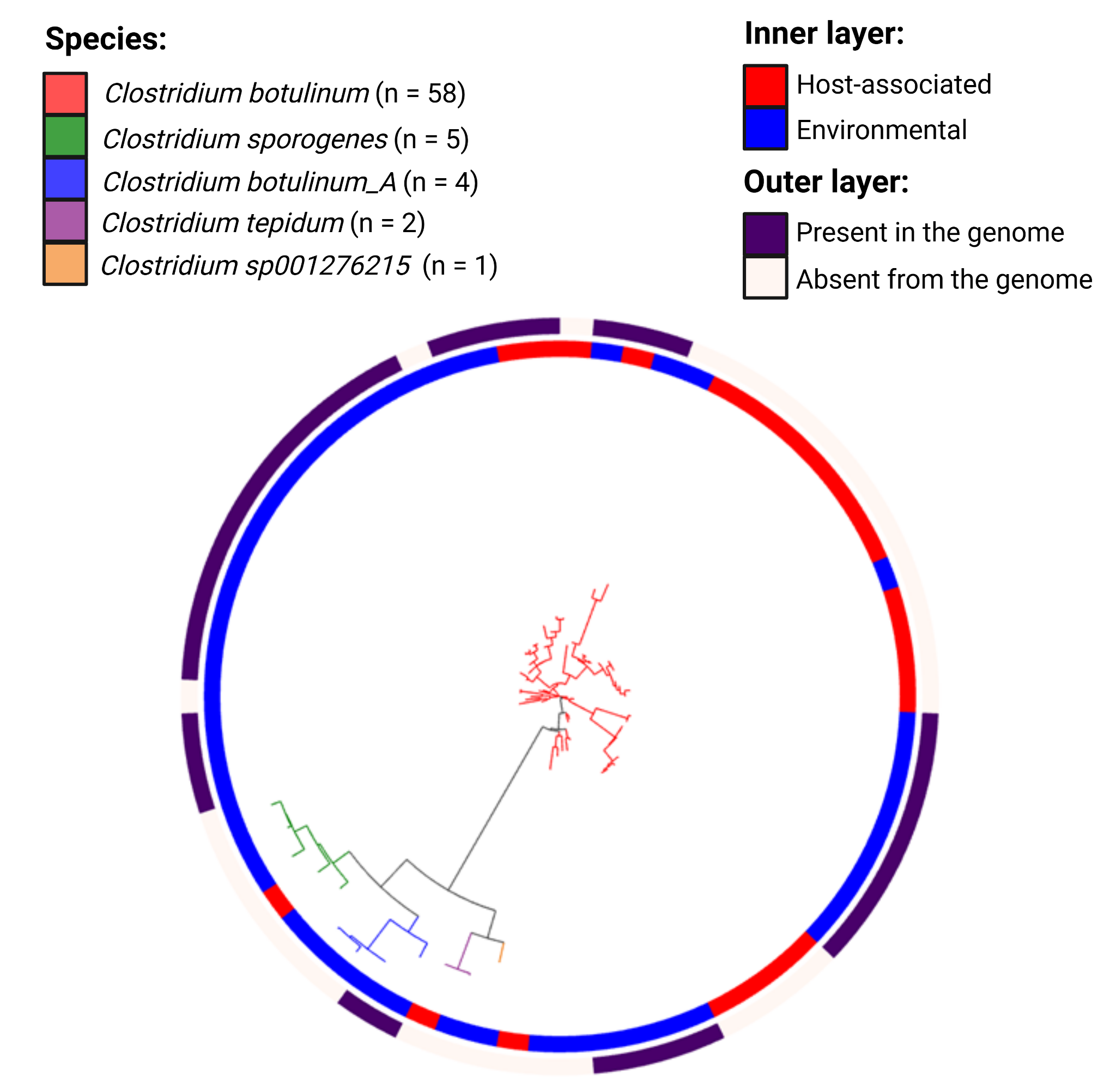

**Supplementary Figure 12: Phylogenetic tree illustrating the presence of Pfam domains associated with the botulinum neurotoxin complex.** The tree is constructed using marker genes from bacterial genomes. Branch colors indicate different species within the genus. The inner circle shows genome labels, with blue representing host-associated and red indicating environmental genomes. The outer circle displays the presence of botulinum neurotoxin complex-associated Pfam domains, where purple indicates presence and white denotes absence.

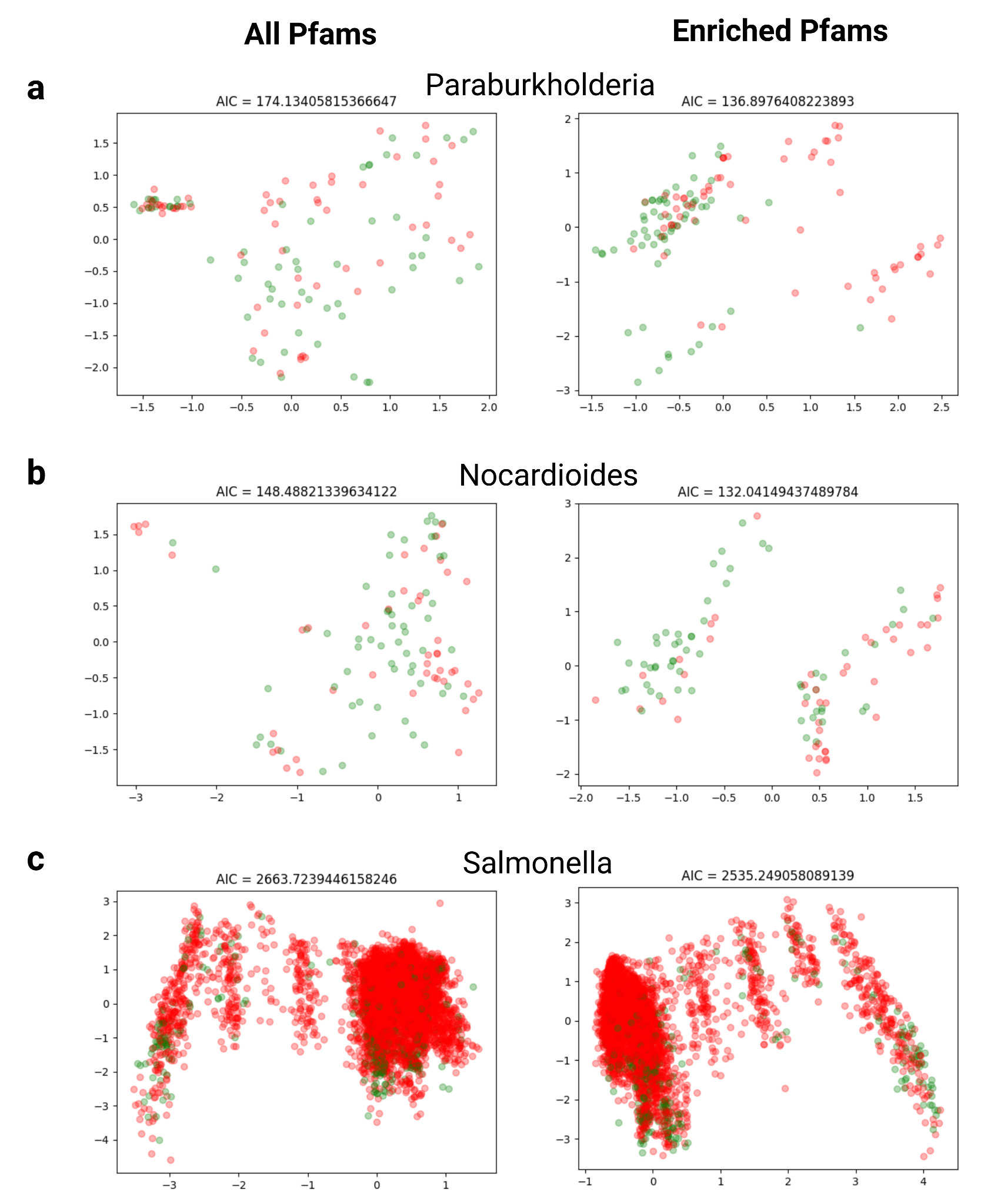

**Supplementary figure 13:** PCoA results of three genera labeled as host-associated in red and environmental in green, using Pfams. On the left side are the genomes clustered by all genes mapped to Pfam, on the right side are the genomes clustered solely by enriched Pfam results. A and B are examples of improvement in the PCoA using the enriched results and C is an example of a clade without significant improvement.

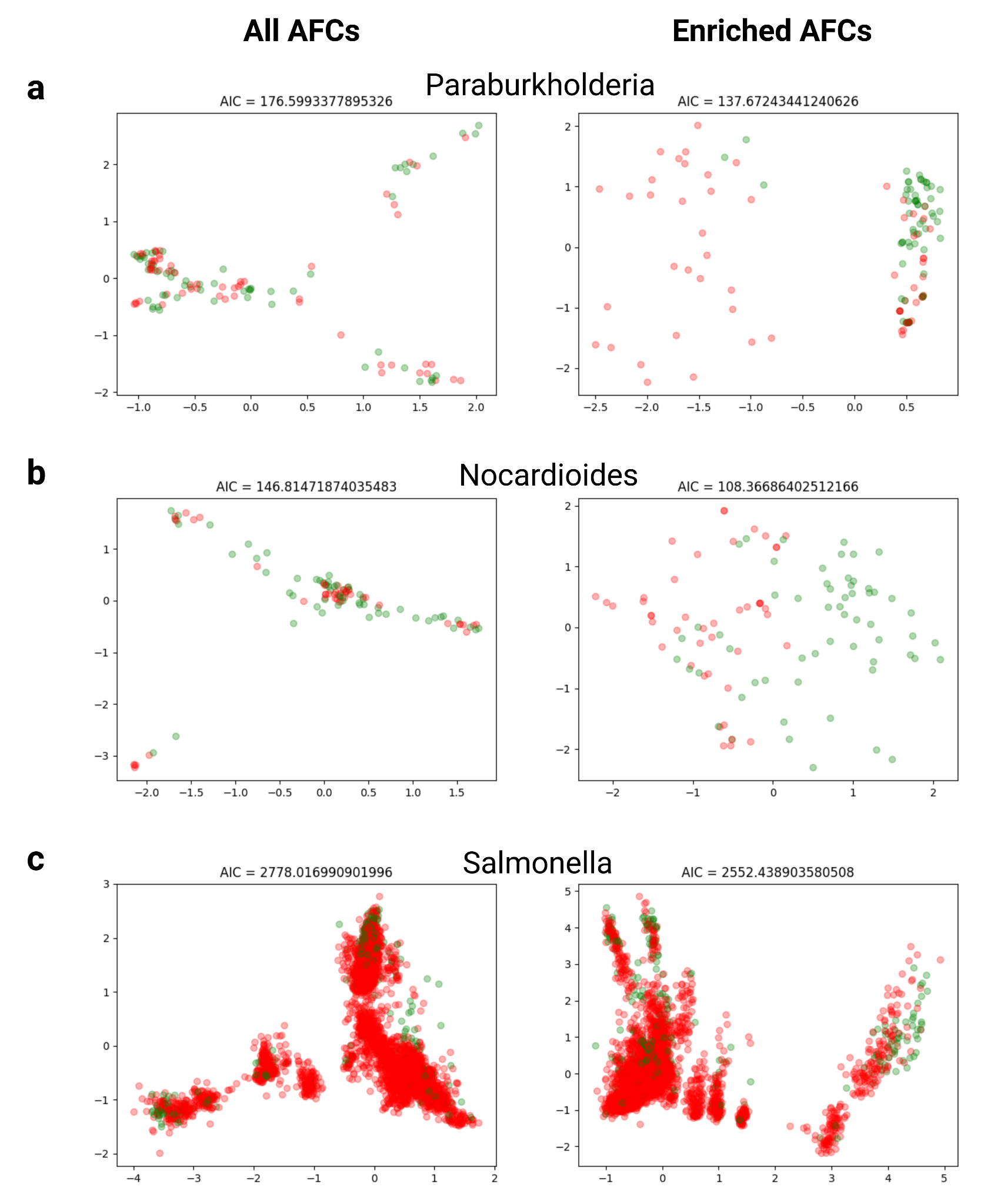

**Supplementary figure 14:** PCoA results of three genera labeled as host-associated in red and environmental in green using AFC. On the left side are the genomes clustered by all genes mapped to AFC, on the right side are the genomes clustered solely by enriched AFC results. A and B are examples of improvement in the PCoA using the enriched results and C is an example of a clade without significant improvement.

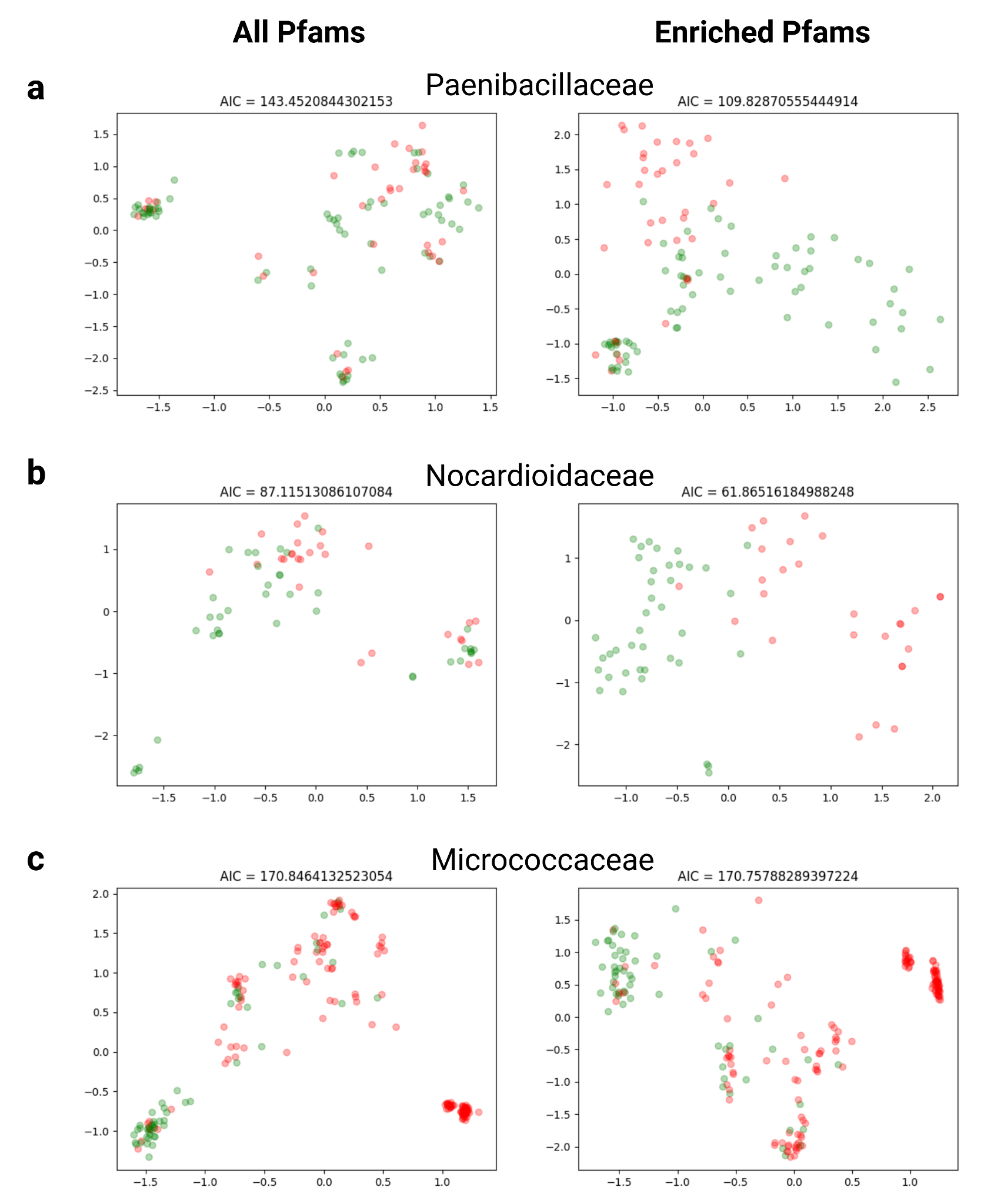

**Supplementary figure 15:** PCoA results of three families labeled as animal-associated in red and plant-associated in green, using Pfams. On the left side are the genomes clustered by all genes mapped to Pfam, on the right side are the genomes clustered solely by enriched Pfam results. A and B are examples of improvement in the PCoA using the enriched results and C is an example of a clade without significant improvement.

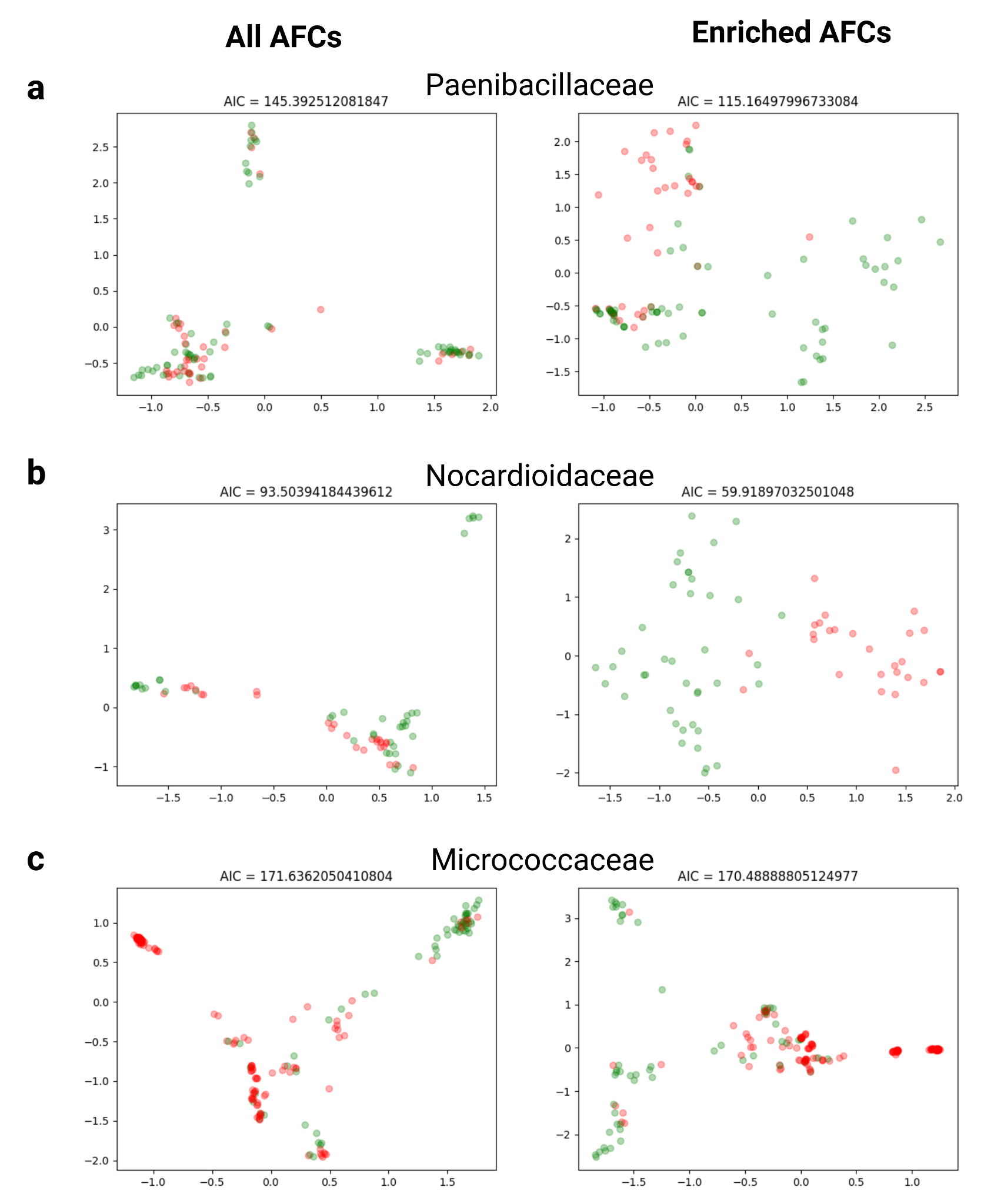

**Supplementary figure 16:** PCoA results of three families labeled as animal-associated in red and plant-associated in green, using AFC. On the left side are the genomes clustered by all genes mapped to AFC, on the right side are the genomes clustered solely by enriched AFC results. A and B are examples of improvement in the PCoA using the enriched results and C is an example of a clade without significant improvement.

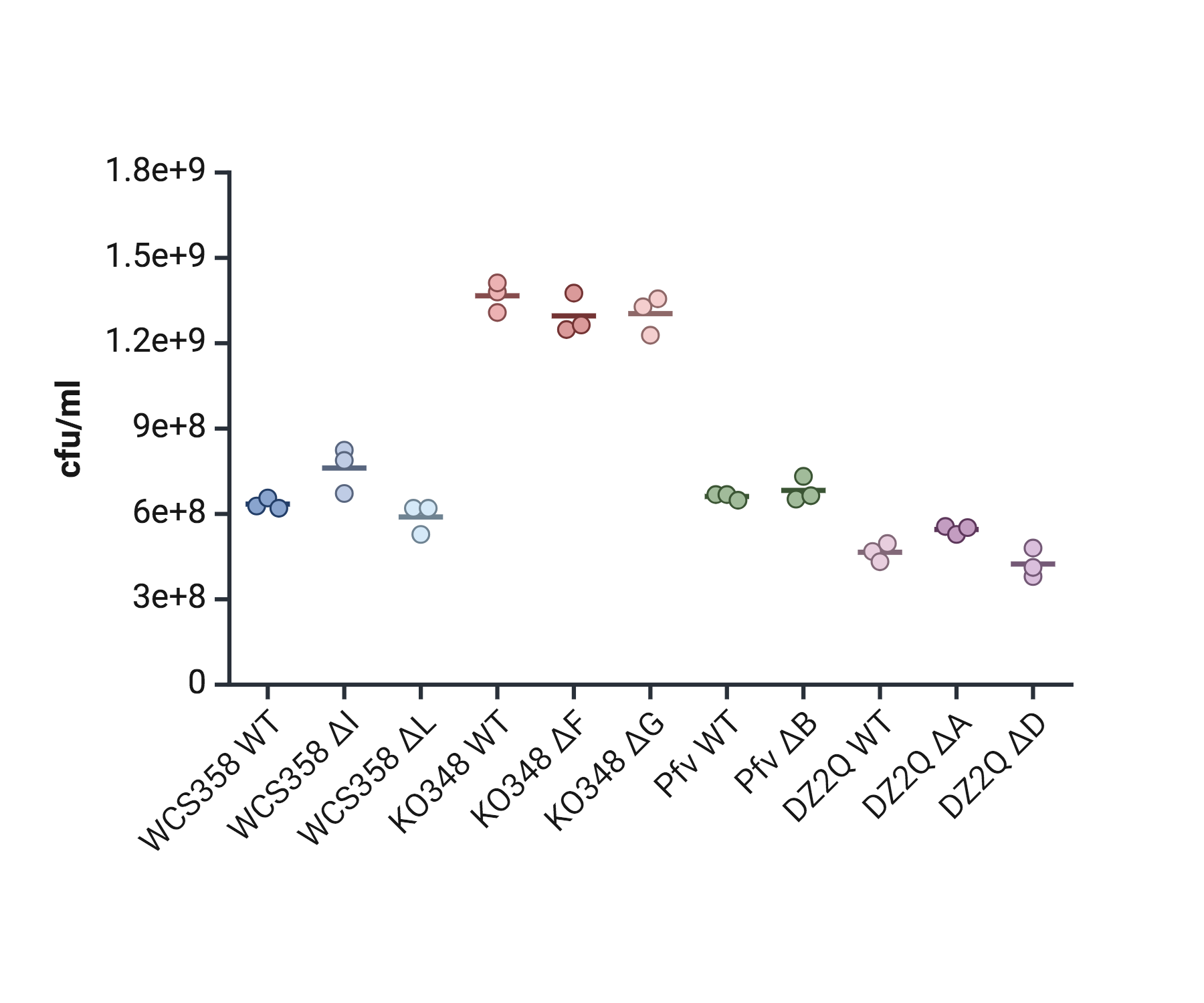

**Supplementary Figure 17: Growth of bacterial strain with their respective mutants.** Colony-forming units (CFU) per milliliter of bacterial strains and their respective mutants grown on Tryptic Soy Agar (TSA) plates after 18 h of incubation. Samples were prepared by performing serial dilutions of bacterial overnight cultures grown in Tryptic Soy Broth (TSB). Data points represent individual biological replicates, and horizontal bars indicate mean CFU/mL values. No significant differences were detected among WT and respective mutants.

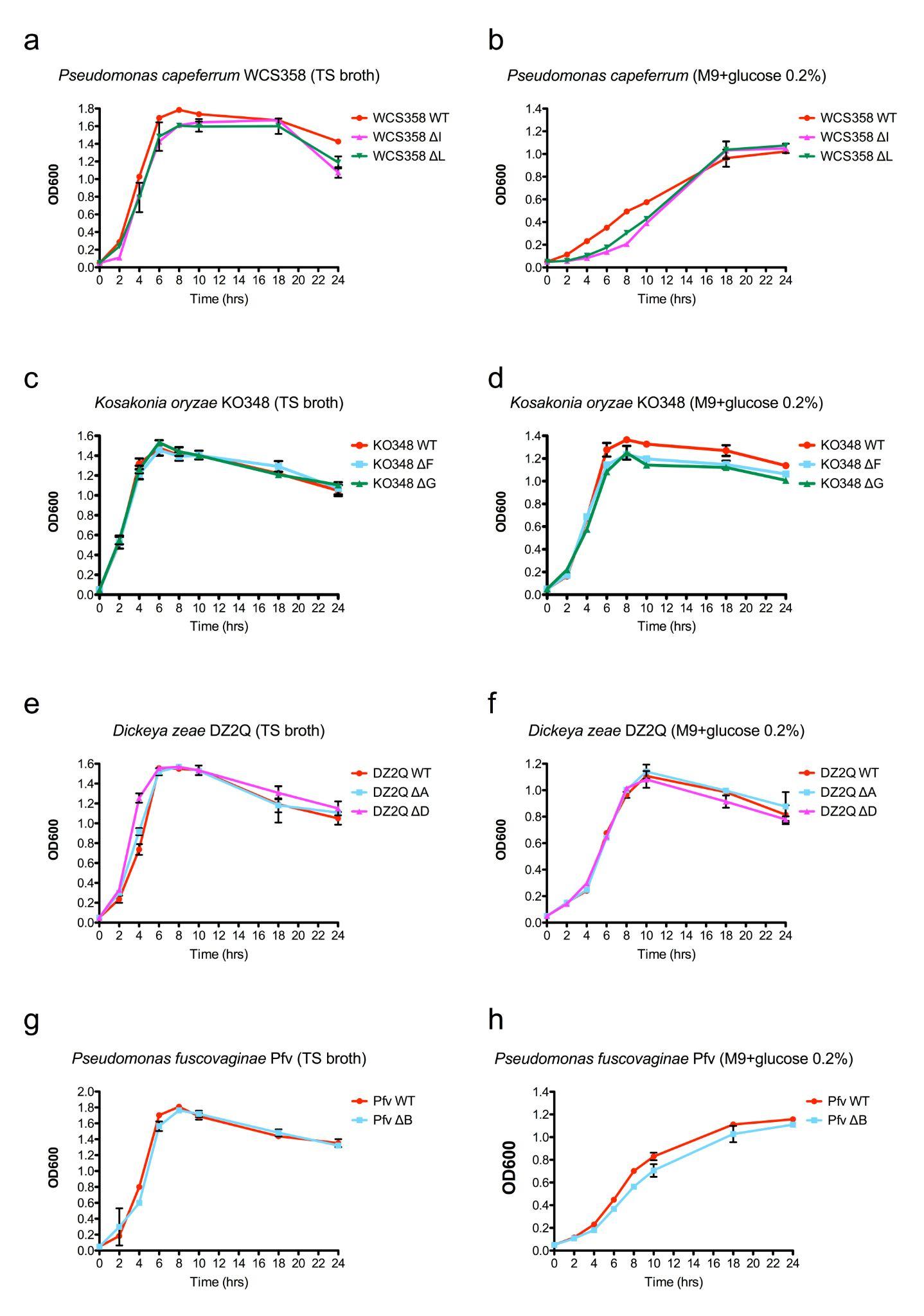

**Supplementary Figure 18: Growth curves of bacterial strains investigated in this study and their respective mutants in two liquid media.** Tryptic Soy Broth (TSB)**,** rich medium, and M9 medium supplemented with glucose (2%) as the sole carbon source. Growth was monitored over a 24-hour period using optical density measurements at optical density (OD) 600 nm. Data represent the mean values ± standard error from five independent experiments. (a-b) Growth curve of *Pseudomonas capeferrum* WCS358 WT, WCS358 ΔI, and WCS358 ΔL in tryptic soy broth and M9, respectively. (c-d) Growth curve of *Kosakonia oryzae* K0348 WT, K0348 ΔF, and K0348 ΔG in tryptic soy broth and M9, respectively. (e-f) Growth curve of *Dickeya zeae* D2Q2 WT, D2Q2 ΔA, and D2Q2 ΔD in tryptic soy broth and M9, respectively. (g-h) Growth curve of *Pseudomonas fuscovaginae* Pfv UPB0736 WT and the mutant Pfv UPB0736 ΔB in tryptic soy broth and M9, respectively.

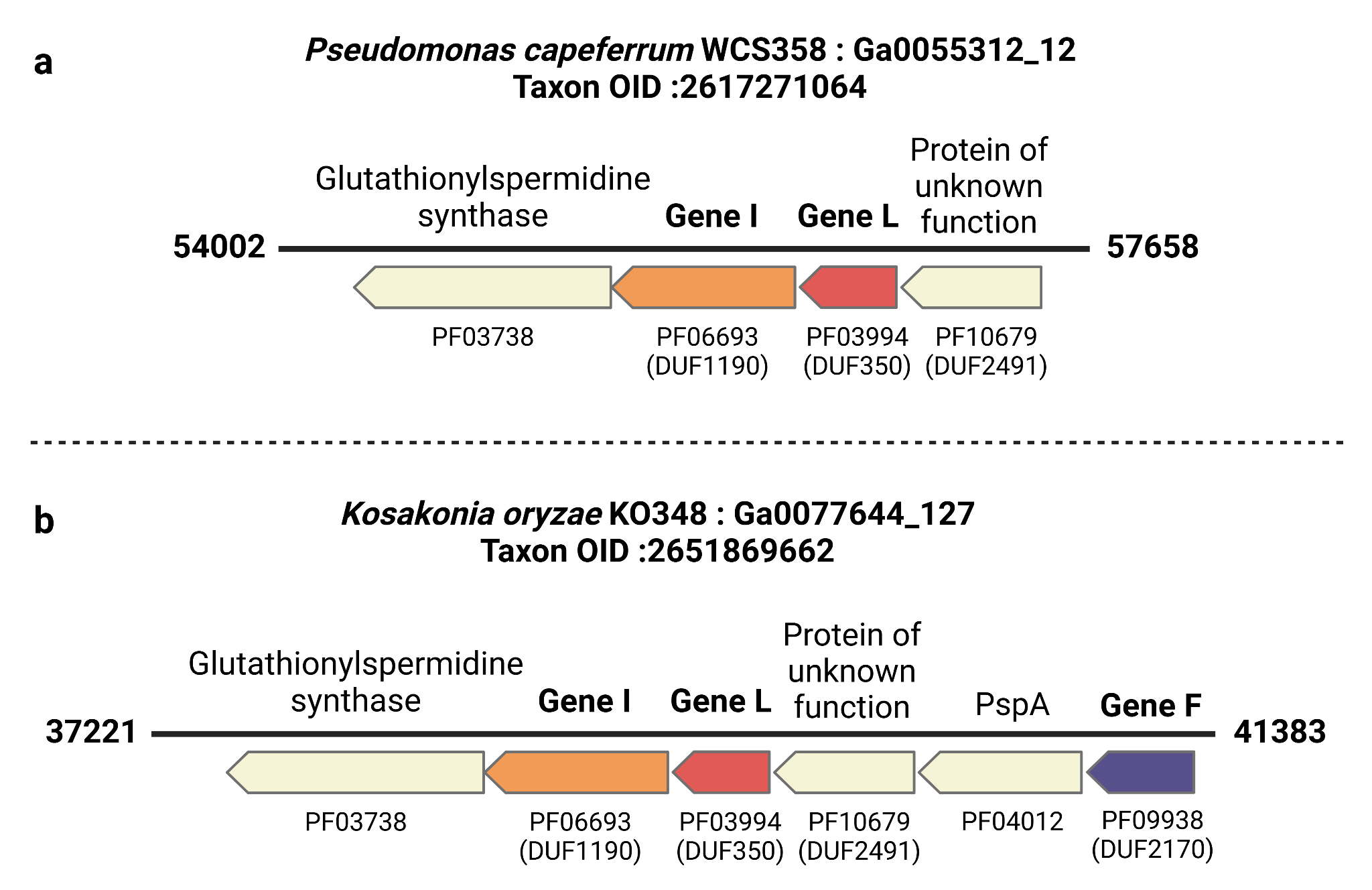

**Supplementary Figure 19: Genomic localization of knockout genes in the root colonization experiment.** Each arrow represents a gene within the genome, with bold numbers on either side indicating its genomic position within the contig. Below each gene name, the corresponding Pfam IDs are listed, as identified within the gene. Colored arrows highlight the knockout genes used in the experimental assays, with the colors matching those in Figure 5 of the main text. Taxon OID refers to the IMG genomes database.

**a, Genomic localization of *Pseudomonas capeferrum* WCS358**

**b, Genomic localization of *Kosakonia oryzae* KO348**

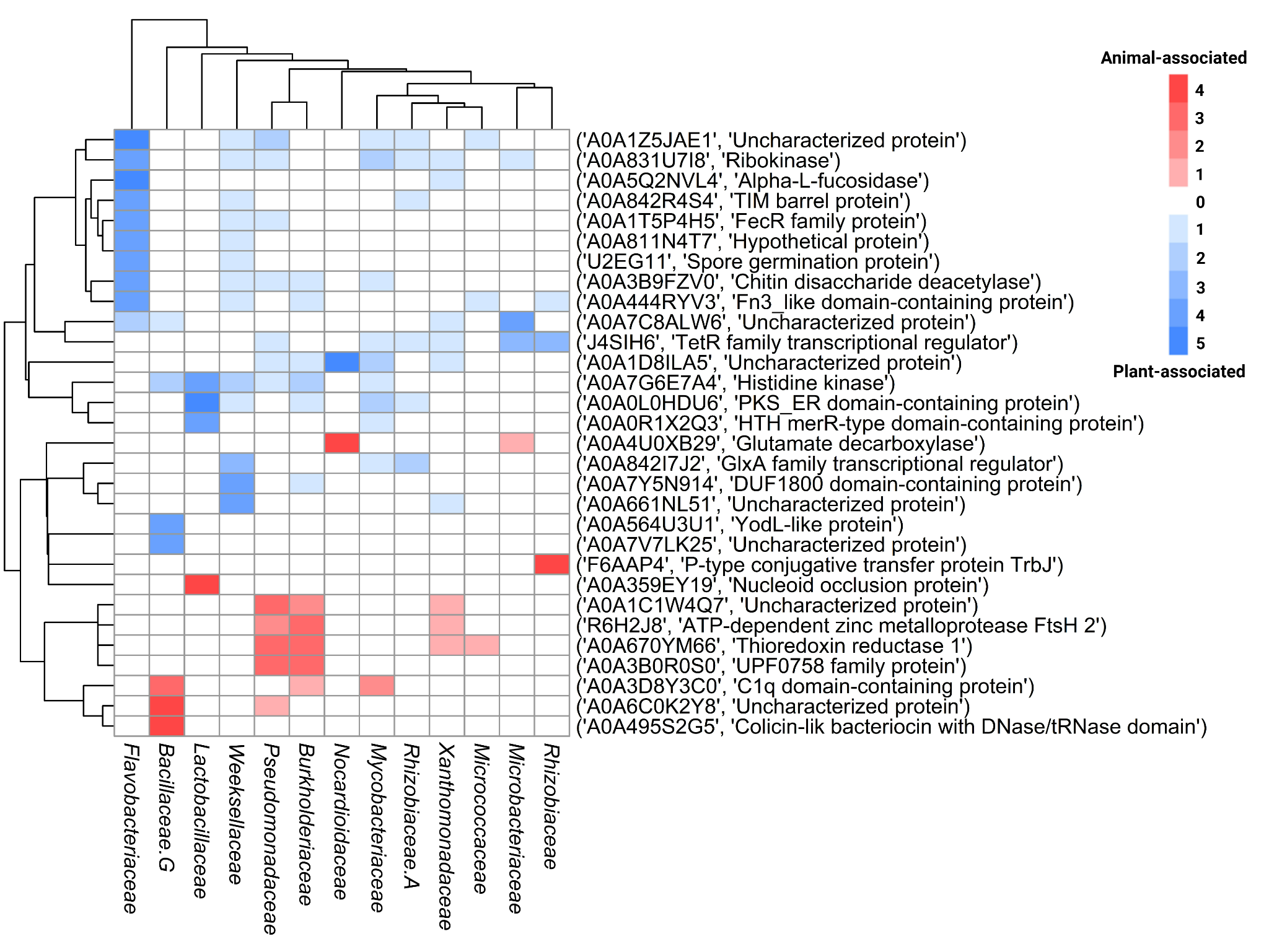

**Supplementary Figure 20: Heatmap showing the enrichment of Animal-associated and Plant-associated AFCs across 13 bacterial families.** Each row represents an AlphaFold cluster (AFC), identified by its UniProt accession and description, while the columns correspond to different families. Red shading indicates the number of tests that identified these AFCs as significantly enriched in animal-associated bacteria within a clade, and blue shading indicates enrichment in plant-associated bacteria. White areas denote no significant enrichment.

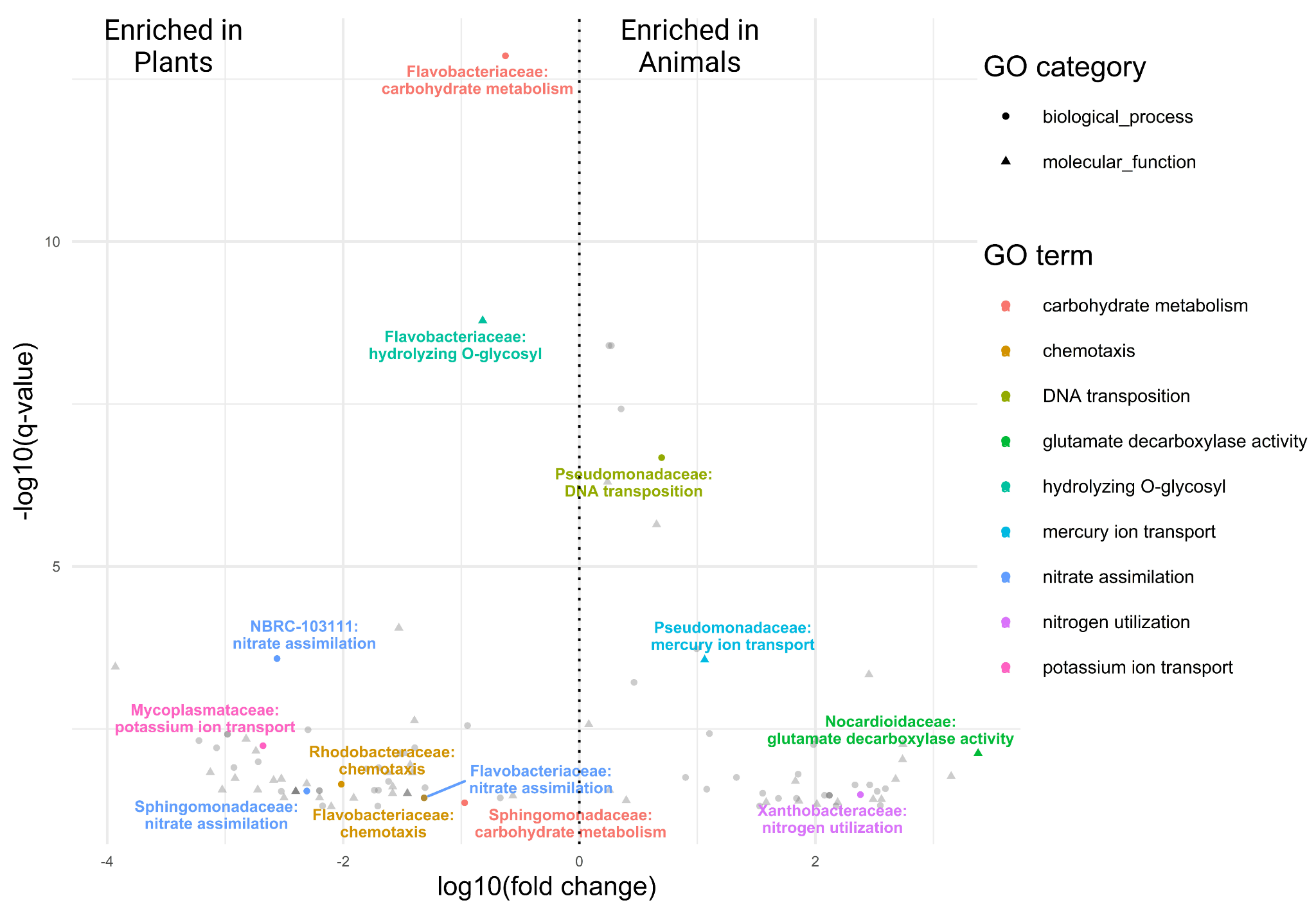

**Supplementary Figure 21: Volcano plot highlighting highly enriched Animal-associated and Plant-associated functions based on AFCs at the family level.** The y-axis represents the significance of enrichment (-log10 FDR-corrected q-value), while the x-axis shows the fold change (log10 fold change) for each mapped GO term. Colors highlight specific key functions. Functions located to the right of the dotted line are enriched in animal-associated bacteria, while those on the left enriched in plant-associated bacteria. To improve readability, "mercury ion transmembrane transporter activity" was abbreviated to "mercury ion transport", "carbohydrate metabolic process" to "carbohydrate metabolism" and "hydrolase activity, hydrolyzing O-glycosyl compounds" to "hydrolyzing O-glycosyl".

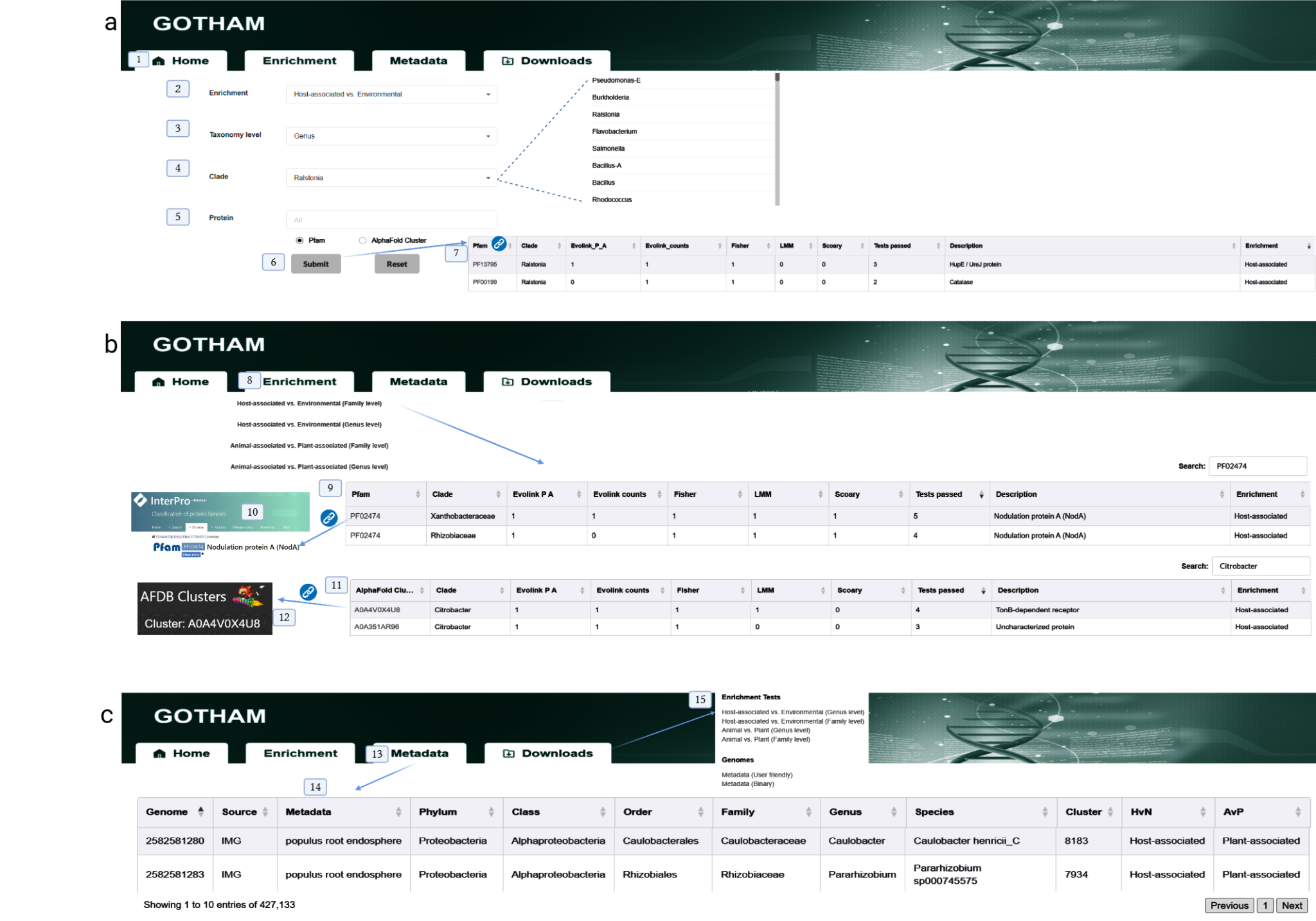

**Supplementary Figure 22: Overview of the GOTHAM Database Interface. A. Home.**The homepage (1) features multiple search parameters that allow users to refine their queries. Users can select comparison types through the enrichment field(2), filter data by taxonomic level (Genus or Family)(3), search for specific clades(4), and input protein identifiers using either Pfam IDs or AFCS numbers(5). Upon submission(6), results are displayed in a comprehensive table format(7).**B. Enrichment.** The Enrichment Analysis section (8) offers specialized comparative analyses through a dropdown menu with four distinct options: Host-associated vs. Environmental and Plant-associated vs. Environmental, each available at both Genus and Family taxonomic levels. Results are presented in two interconnected tables - a Pfam table(9) providing direct links to InterPro entries for protein families(11), and an AFCS table(10) linking to corresponding entries in the AFDB Cluster database(12)**.C Metadata and Downloads.** The Metadata tab(14) provides detailed information for all database entries, while the Downloads section(15) enables users to access complete datasets in various formats (16).

**Supplementary Table 14. Primers and plasmids used in this study.** List of plasmids and primers used to generate the knock-outs and to check them via PCR.

| **Primers** | **Sequence (5’-3’)** | **Reference or sources** |
| --- | --- | --- |
| FW_*geneA*_ext | ATG CTG CTG TCA GCT GGC TT | This study-mutant check |
| RV_*geneA*_ext | AAC GCG GCA CTG CGA CAT G | This study-mutant check |
| FW_*geneB*_ext | CTG CCC AAG GCC GCA GAA C | This study-mutant check |
| RV_*geneB*_ext | TCG TCC GTC GCC TGC TGC A | This study-mutant check |
| FW_*geneD*_ext | ATC AGA GGT CTG CGG AGC ACA T | This study-mutant check |
| RV_*geneD*_ext | TCT TGC CAT TTA CGG GTA CGC A | This study-mutant check |
| FW_*geneF*_ext | TGC ATG CGG TGA TTG TGC CGA | This study-mutant check |
| RV_*geneF*_ext | CTG TTG CGC GCG CTG CAT C | This study-mutant check |
| FW_*geneG*_ext | TGG CGC GGG AGG AGA ACA C | This study-mutant check |
| RV_*geneG*_ext | CAG TGT TCC GGC GTG TCG TG | This study-mutant check |
| FW_*geneI*_ext | TAC TCC GCA CCG CGA GTT CG | This study-mutant check |
| RV_*geneI*_ext | TAC AGC GAG GTA GGC GTG TC | This study-mutant check |
| FW_*geneL*_ext | CTC GGT GCT TCG ATG CAC CTG | This study-mutant check |
| RV_*geneL*_ext | GCG CCG TCG CGA TAG GTA | This study-mutant check |
| FW_comple_geneI_Bam | TTGGATCCTCAATTCGGCTTGCATGACC | This study-mutant check |
| Rv_comple_geneI_Hind | TTAAGCTTGGGCCTGGCTGAATTGG | This study-mutant check |
| FW_comple_geneL_Eco | TTGAATTCCGA GCA GGG CAA CGC CAC TT | This study-mutant check |
| Rv_comple_geneL_Hind | TTAAGCTTCAA AGG CAG CGA GCT GGC | This study-mutant check |
| Plasmids | Relevant features | Reference or sources |
| pEX19Gm | Suicide vector for making deletion mutants, Gm^R^ | Dreier J, Ruggerone P. 2015. Interaction of antibacterial compounds with RND efflux pumps in *Pseudomonas aeruginosa*. Frontiers in microbiology 6:660 |
| pUC4K | pUC7 derivative, Amp^R^ and Km^R^ | Addgene, Watertown, MA |
| pUC57 | Cloning vector; Amp^R^ | Addgene, Watertown, MA |
| pLAFR3 | Broad-host-range vector; IncP;Tc^R^ | Staskawicz B, Dahlbeck D, Keen N, Napoli C. 1987. Molecular characterization of cloned avirulence genes from race 0 and race 1 of *Pseudomonas syringae pv. glycinea*. Journal of Bacteriology 169:5789-5794 |
| pPHJ1J | Broad-host-range vector; IncP;Gm^R^ | Addgene, Watertown, MA |
