## Supplementary Dataset 1 for "The Genetic Basis of Bacterial Adaptation to Hosts"

**Supplementary Dataset 1:** List of constructs synthesized synthetically and subsequently cloned into the suicide plasmid pEX to generate the respective knock-outs. Each sequence consists of two 600-nt sequences joined with a central deletion of 200 nt from the gene, thereby creating a chimeric sequence.

**>geneA_Dickeya_PF07511**

**AAGGTACCtgccccactcttcaagaatgaagggttttatgaggaatctgagtggcgcatcatagctagggggccgcgagagcctgtgcattttcggccaagttcctcctatttggcaccatacgtcgagttaaaagttctttccgatactgcggtgtctgctttgcgggaagttattgtcgggcctaatcccaatcagcgtcgttgtgaatcatctatcaagatgctcctggattcttgtgggttaggcggtgtcgaggtaaggatgtcttccttaccatttaatagctggtgatccagggatcgactttatccgttctttgttacgggacataccctagtctggctctgacgacaggaagtcgattggttaattttttgaatctcagaaaacagttccttactgacacgccgtgcccataaacgcaccttaaacgggttagttttccccgtttaaggagtgctcaccATGTTCAGATTTGTTTATCTGACGGCGCTGTTGCTGCCTGCTGGTACGTTTGCCGGTACGGTCATTTATACCGACAGTGCTCATCCCGTTATACCTGGCTCATCCGCCGCCCAGGTCGTCTACCTCGATGCTCCCGATAAggatccCGCCGGCGTGATCAAGGCCTGGCAACTGCAGCTGCAGCGTTATCCCGCTGTCGTCTTTGATGATACCTATGTTGTCTATGGTACTACCGACGTCGATCTGGCCAGTACCAGGCTGAGCAACTGGCAGGAGGCGCGCCGATGAggatgacgctgaaaacccgacgggtgctgatggccaccctgttatctgtcagcgggctgacaacggccagcgtaaatacggcgcagattgtggccagcgcatcgtcggtatcgtgcatcagttggcgagtcaagggtatctgctactggctgctctgtaccccatttggctgctcggttaagacctcggtccgcgtcgagcatttcattccagaggctgtcgtatcggcgtacggtaacagcggtgataacccctggactgaaatgtcgctggtcagccagagcgcagcatctgcagaaggtggattgatcgggagtatcgccggcgttgttgccggcgggggtaatcaggaactgaaagcccccgaagcagggcgtaagaagaacttgcgttatctctatagcgatgcgattggtcacccggctacgtcattgattggtggaatggtcccgggttacAAGCTTAA**

**>geneB_Fusco_PF07511**

**AAGAATTCggtggatttgcctgaacatgccgccggatgaggtcaacaagatcgcccgcttccgctcgctcaccgacgcgcagaaaaccctgatgctgtccgcccgcaaggagtcggggaaattcaccgagggggtaatcctgtccaagtccatggaggtgctgtttcgcgcggtaccgccaagcctctacctgaccatggcgatgacggagccggaggaaaaggccgaacgctttgcactgatgaatgcccggggcatcagcgaactggaggcagccatggccgtggcccaacggatggatcaggcgcgcggcatcactacgctggcagaaggtgcggcctgcatcgattaccgtgacgaagaggcagagccgccgcgataaaagcggcaacccgcaaaacagttttccggctgtgatcacccaggtttttcgcgaggattcggttttctgccaaccggaccctcgccATGACGAACCCTCTGCTCCTGCGACTGCTGCGACTGTGTAGCCCCGCGACCCTGCTTGCCTGCCTATCTCCCTGCCTGGCTGAAACCTGGGTCATCACCGATTCGGCACACCCCGTCGACCTGCctgcagATACGCAACTCGATCGCGCTCATCAAGACATCGTCAATGCCTGGAGTCTGGGGGTCAGCAAGATTCCGGCCGTCGTCGTCGACCGGCAGTACGTCGTCTACGGCGACCCGAACGTTGCGCGTGCGCTGACGAAAATCGAGCATTTTCGTCGGGCCAGCCGATAAgggcctgcacatgtcgggtcgccggggacctaccgatggattcagtcgttggcccgcaaccgtgcgaccgggtgctcatcgaccccgggcacggcggcaggccacgagccctccagcgcaagcccgtcagccgccacgccgagtcgactcccgatcagcggcggcgcgaaatcggcacggcggctgccctccagcacgtccaatggcaactcgccaaacagcgccctggcctggcgagcctggtcgttatcggcggcgaaatcggcccgcgtcacccccgaactgcggtactggcgaggcaacccgaacaggccgcgcaactggcgagcgccggcatggatccggctttccatctcccgccgatcaatcaacgccacgtgatgagcAAGCTTAA**

**>geneD_Dickeya_PF12101**

**AAGAATTCaaataacggcgcaggtaggtgactactctgcgccgttatttttttcggcggatgaataattatccatccccactactgttgacttaacttgttcccggtttaatgattaaggcgaaccttcgttgtctttgacaacagccaatgttccagtggtttgagaggaacgccttcgggtaatcacgataaatgaagatcatccgcataaccctcataagcgagttatgtgacctggaagatgatcgccttaacaggtggtaaagccgtttaacctttgagaacggcgaatgcacaacagcatttaaggtcatttcaacttagcagtcagtctaagcgcttagtctgacttccctgcgggaaacatttttcccgtcgggaatgaatgtttttccgcatgacatctttttactccggagaaacatcATGACTGCATCAGCAAAATCAACCACAGCAACTGAAACTCAGTACTTTAATCTCAACACCGCAGGTATCGGTTATCTGTCCAATATTCGTCAGGTTTCTGGCGATAAAGGCCCTTTCACCTGTGCTGTGATTAACGCCTTGTCTGGTCCAACGGACAACGCGTCGTATATACGTTTTGACgtcgacCTGAAAAGTGGTGAACACGCTGGTGAAGATCGCATCGCGCTTCGCTCCCGTCTGATCAAGATCGACTTTATCAAAATCGGGAAAGAGACGGTTTATAAGGCTGAAAAGCCGACGTCTACAACACCGCCGGCACAGAACGAAGCTCAGCAGCAGAAACAATACGCTGCTAATTCCTTCTGAaccgattcggaatcacaccatcaggaacatgcctgatggtgtgttcctcttcaaattttgacttattgaggaatgaatcatgcacggcgttaacaaaattattctgctcggtttcctggcgatagccttgatgccgaaaggattttttaatctgccagtggtttttttcttgctgatgtttgttcctcttcaaacttttattgaggaatgaatcatgtctgcacgcggtgttaacaaagtaatcctgattggcttcctgggacaggatccggaagttcggtctatgccaaacgggactgcagtaacaggatttcagattgcaacgtcagaaacctggcgtgacaagcaaagcggggaacagaaagagcggaccgaatggcatcgtatctctctgtacggcaaattggctgagattgctggAAGCTTAA**

**>geneF_Kosakonia_PF09938**

**AAGAATTCGAATGCCACGGCGAAAAAAGTCGCCTGCGGCGCGGCAGAGAGCGTACCGTTGATCCGCGTGACCAACCTGGCGCGCACCATGCGCCTGTTGCAGGAAGAGAATATCTGGATTGTCGGCACAGCGGGCGAAGCCGATCACACCCTGTTCCAGAGCAAAATGACCGGGCGCATGGCGCTGGTAATGGGCGCGGAAGGTGAAGGTATGCGCCGCCTGACGCGTGAGCACTGTGATGAGCTGATCAGCATTCCGATGGCGGGCAGCGTGTCGTCGCTCAACGTCTCGGTCGCCACCGGCATCTGCTTGTTTGAAGCGGTGCGCCAGCGCGCGCTTTAAACTTCCAGCGCACGGTAAAACCACCCTTATTCGGGTGGTTTTTTTTGCCCTGAACACTGGTAGAATTTGCCGGCAGATATTTTCTCTTTTTACTTTTATTGCGATGGAATAATCAGGGTATAAATGatggcatggactgtagaaacacttggcgcggcattattgcaatcatcgacgcttacgttaaatatccaacacgaagctgatgcgttaattattaaactggatgactacggcgatttgcaattaaatctgttgctcacttcGGATCCgcctgtcggcggggaagagtattatgtcgccttcggcgcgctgtcgcttaactcctcgctcgacgatatcctgctggaaatctccacgcttgcgcaaaacgcgctggatcttgctgaactaaccgatgattttacccaataaCATTGTCAGGGAGTGGTTGTATGGGAATTTTAAAAAGCCTGTTTACGCTGGGTAAATCCTTTATTGCTCAGGCGGAAGAGTCGATTGAAGAGACGCAGGGCGTGCGCATGCTGGAGCAACACATTCGCGACGCGCGCGCTGAGCTGGATAAAGCGGGTAAATCCCGCGTCGACCTGCTGGCGCGCGTTAAGTTGAGCAACGACAAACTCAACGATTTACGTGAGCGCAAAGCCAGCCTTGAAGCCCGCGCGCTGGAAGCGATGGCGAAAAATGTGGATGCCGGTTTGCTGAATGAAGTCGCCAGCGAAATCGCCCGTCTGGAAAATACCATTGCGGCGGAAGAACAAGTGCTGGCGAATCTTGAAACGTCGCGCGATGCCGTTGAAAAAGCGGTAACCGCCACGGCGCAACGTATTGAGCAGTTTGAGCAGCAACTGGAAGTGATTAAAGCCACCGATAAGCTTAA**

**>geneG_Kosakonia_PF11844**

**AAGAATTCATGAGCCAGACTGCTGAGAAAATACAGAATTGCCATCCCCTGTTTGAACAGGATGCTTACCAGACGTTATTTGCCGGTAAACGGGCACTCGAAGAGGCGCACTCGCCGGAGCGGGTGCAGGAAGTTTTTCAATGGACCACCACCCCGGAATACGAAGCGCTGAACTTCAAACGCGAAGCGCTGACTATCGACCCGGCAAAAGCCTGCCAGCCGCTGGGTGCAGTGCTCTGTTCGCTGGGGTTTGCCAACACCCTGCCATATGTGCACGGTTCACAGGGTTGCGTGGCCTATTTCCGTACGTACTTTAACCGCCACTTCAAAGAACCGGTGGCCTGCGTGTCGGATTCAATGACGGAAGACGCGGCCGTGTTCGGCGGGAATAACAATCTCAATACCGGGCTACAAAACGCCAGCGCGCTGTATAAACCGGAGATTATCGCCGTCTCTACCACCTGTATGGCGGAGGTAATCGGCGATGACTTGCAGGCTTTTATCGCCAACGCTAAAAAAGATGGTTTTCTTGATGCCGCCATCCCCGTGCCCTACGCGCACACCCCCAGTTTTATCGGCAGCCATATCACTGTTCCCGTTGGGCTGGCggatccAGGAACGGATGAACTATTAATGGCGATCAGTCAGTTAACCGGCAAGGCCATTCCCGATTCACTGGCGCTGGAGCGCGGGCGGCTGGTCGATATGATGCTCGACTCCCACACCTGGCTGCACGGTAAAAAATTCGGTCTGTTTGGCGACCCGGATTTTGTCATGGGATTGACCCGTTTCCTGCTGGAGCTGGGCTGCGAACCGACCGTTATCCTCTGCCATAACGGTAACAAGCGCTGGCAGAAAGCGATGAAGAAAATGCTCGACGCCTCACCGTACGGGCAGGAGAGCGAAGTATTTATCAACTGCGATTTGTGGCATTTTCGCTCGCTGATGTTTACCCGCCAGCCGGACTTTATGATTGGCAACTCGTACGGCAAGTTTATTCAGCGCGACACCTTAGCCAAGGGTGAACAGTTTGAAGTTCCGCTGATCCGCCTCGGTTTTCCCCTGTTCGACCGCCACCATCTGCACCGCCAGACCACCTGGGGTTATGAGGGCGCCATGAGCATTCTCACTACCCTTGTGAATGCGGTACTGGAGAAAGTGGACAAAGAGACCATCAAGCTCGGCAAAACCGACTACAGCTTCGATCTTATCCGTTAAAGCTTAA**

**>geneI_WCS358_PF06693**

**AAGGTACCTTCGCTGCCGGCCAGCAATATCATTGCCTACAGCGTCGGCATCCTCGACGTAGTGGCTTGGGTAGTGATTGCGGCAGTGGTGCAGTTGCTTGCCTTCGGCCTTACCAGCCTGGTGCTGCGCGGCCTGTCCAGGCGTATCGCCGCAGGCGAACTGGCCGCTGCCATCTACGCTGCCAGCGTGGCGATCAGCGTCGGCTTACTCAATTCGGCTTGCATGACCCCGTCGGCCTGATGCACTAGGAGCCCCCatgaaatccccaatgagacgaagctccgtcaagctggtgctggccagctcgctgcctttggcgttgaccgcgtgcagcccgcaagaagaaacctacaccgtcagccagcaggtgaactacggcagtgtcgaagcctgcgtcaacgacaaggtgccggagaaggcgtgcaacgatgcctacaagcaggcgctggccgagtaccggcgcaccgccccgacctatgtatcgaagcccgactgcgaagccgaattcgcctctgacggttgtcagataggcgccggtggccgttacatgccccgcatgagcgggttcgagctggacaccgcaggcgaagtcacccagtcccaggtcgGGATCCcctatcgcgacggcgccgggcagcgcagtggcctgcttgggcagcgttatggtgtggccgggggcattcagcgcaccaacaaggacgagcagagcagcagtggcggttcgtccggcggcggcggtgctggctatggcgcctcaggccgcacggcgtcgcgctcggccgtctctgcgtccatctcgcggggtggctttggcagccaggccacggcccgcagtggctggggcggcaagtctggcagcttcttcggggggtagGCATGCGCAAGGTCGAGCTCGCAGAGCGCCCAGGCTGGCGCGCCACTGCCCAACAAGAGGGCTTTGCCTTCCACACCCTCGATGGCGAGCGCTACTGGGACGAGCGGGGCTACTACCAATTCAGCCAGGCCCAGGTCGAGCGCGACTTGCAGGCCCCGACCGAAGCACTGCACGCGATGTGCCTGGACGCCGTTGATCGTATCGTCGACAGCGAGGCGCTGATGACGCGCCTGGCCATTCCTGCGGCATTCTTCGACCTGGTGCGGCACTCGTGGCTAGCGCGGCAGCCGCATCTGTATGGGCGCTTTGACTTCAGCTATGACGGCCATGGCCCGGCGAAAAGCTTAA**

**>geneL_WCS358_PF03994**

**AAGGTACCGAAGACTATTGGCTGCAAGTGCTGATGAACGGTAGCGAAGTGGGTGATACCGTATTGTTCGGTTATCACAGCGCTGTACCGATCCGCGACACGTCGCACCTTCAGCGCCTTGTCGGCAGCCAGTCCAAGGTCGGCCTGCCGATCTACGAACACGACGACTACCTGTATTCCCGCCAGTGGGGCCAGGAGGCAGGCCAGGCCAAACTGGTTCCGTTGCACGAGCGCGTGACCAGCCCTGATGCGCGCTATGGCATCCAGCACCTGTCGATGCTCTACGCCCGCGATACCGGCCTGCCAGAGCGTCGCGAATTCCTGCTGCTGTCAGCGGAGGAGGACGAGCAGGGCAACGCCACTTTCACCACTTCGTTGGGCGTCACCCTGCACCCCACGGACTTTCACGTAACCTGACAAGGACTCACCatgctcgacgctctgcgcctgtcgctcaatgccactgcagtgactggcttcatcctttatatcagcgtggcactgttgctgttctggctgttccagttcatctatacccgccttactccgcaccgcgagttcgcgctcattcgtgagaacaacccggccgcggccattgccctgggcggcGGATCCagcctggtgctgcgcggcctgtccaggcgtatcgccgcaggcgaactggccgctgccatctacgctgccagcgtggcgatcagcgtcggcttactcaattcggcttgcatgaccccgtcggcctgaTGCACTAGGAGCCCCCATGAAATCCCCAATGAGACGAAGCTCCGTCAAGCTGGTGCTGGCCAGCTCGCTGCCTTTGGCGTTGACCGCGTGCAGCCCGCAAGAAGAAACCTACACCGTCAGCCAGCAGGTGAACTACGGCAGTGTCGAAGCCTGCGTCAACGACAAGGTGCCGGAGAAGGCGTGCAACGATGCCTACAAGCAGGCGCTGGCCGAGTACCGGCGCACCGCCCCGACCTATGTATCGAAGCCCGACTGCGAAGCCGAATTCGCCTCTGACGGTTGTCAGATAGGCGCCGGTGGCCGTTACATGCCCCGCATGAGCGGGTTCGAGCTGGACACCGCAGGCGAAGTCACCCAGTCCCAGGTCGATGCGGCCCACGCTCAGGGCGGCGGCTATGGCCACGTCGCCACCGCCGTGGTCGCTGGCATGCTGCTGGGGCAGATGACCAACAGCGCCGACCGCCGTTATCGCGCAAGCTTA**
